# Phage-Assisted, Active Site-Directed Ligand Evolution with a Genetically Encoded *N*^*ε*^-Butyryl-L-Lysine to Identify a Cellularly Potent and Selective Inhibitor for the ENL YEATS Domain as an Anti-Leukemia Agent

**DOI:** 10.1101/2023.01.08.523168

**Authors:** Peng-Hsun Chase Chen, Xuejiao Shirley Guo, Hanyuan Eric Zhang, Zhi Zachary Geng, Gopal K. Dubey, Carol A. Fierke, Shiqing Xu, Wenshe Ray Liu

## Abstract

Eleven-nineteen leukemia protein (ENL) plays pivotal roles in the leukemogenesis. As a YEATS domain protein, ENL reads histone acylation marks and recruits key transcription factors to leukemic drivers such as *HOXA9, MEIS1*, and *MYB* and therefore promotes leukemia development. The histone-reading function of ENL has been proven essential in the onset and progression of several acute leukemias, suggesting a putative therapeutic window for ENL inhibition. In this study, we developed a phage-assisted, active site-directed ligand evolution (PADLE) approach for the identification of potent and selective ENL inhibitors, where *N*^ε^-butyryl-l-lysine (BuK) that possesses known target-protein interactions with the ENL YEATS domain was genetically incorporated into a phage display library to serve as a warhead to direct displayed peptides to the active site of ENL YEATS for enrichment. Using this novel strategy in combination with structure-activity relationship that replaced BuK with other ncAAs for de novo π-π-π stacking interactions with two aromatic residues in ENL YEATS, selective and potent ENL inhibitors with a *K*_*d*_ value as low as 2.0 nM were identified. One pentapeptide inhibitor **tENL-S1f** displayed selective inhibition of ENL over other YEATS domains as well as strong cellular target engagement and on-target effects in inhibiting leukemia cell growth and suppressing the expression of ENL target genes. As the first of its kind study, the current work opens a large avenue of research of using PADLE to develop selective and potent peptidyl inhibitors for a large variety of epigenetic reader proteins.

## Introduction

YEATS domains are newly discovered epigenetic readers involved in many critical transcriptional regulations.^1-2^ In humans, four YEATS domain-containing proteins ENL, AF9, YEATS2, and GAS41 bind to *N*^*ε*^-acetyl-L-lysine (AcK, lysine acetylation) in chromatin and form complexes with chromatin-associated proteins that take part in transcription elongation, histone modification, and chromatin remodeling.^2-3^ Dysregulations of YEATS proteins have been linked to the onset and progression of cancers.^4-7^ In *mixed-lineage leukemia* rearranged leukemias (MLL-r leukemias), *MLL1* (MLL) located on chromosome 11q23 is translocated and fused with partner genes including *MLLT1* (ENL) and *MLLT3* (AF9).^8-10^ The resultant chimeric proteins MLL-ENL and MLL-AF9 bind to several multisubunit complexes involved in transcriptional activation through a shared, highly conserved region ANC1 Homology Domain (AHD).^11-13^ A common mechanism of leukemogenesis is mediated by the histone H3K79 methyltransferase DOT1L. By binding to AHD of MLL-AF9/ENL, DOT1L is localized to MLL1 target genes, such as *HOXA9* and *MEIS1*, resulting in a hypermethylation of H3K79 at these loci and subsequent oncogene expression.^13-17^ In addition to DOT1L, the AF4/FMR2 family members (AFF1-4) bind to AF9/ENL AHD and form the super elongation complex (SEC).^12-13, 18^ During the process, AFF4 recruits a kinase, the positive transcription elongation factor b (P-TEFb) heterodimer (CDK9/cyclin-T1), which activates leukemic gene expression (e.g., *HOXA9, MEIS1, and MYC*) through phosphorylation on the carboxy-terminal domain (CTD) of RNA Polymerase II.^12-13, 19-22^ Together, the co-localization of two transcriptional complexes, DOT1L and SEC, mediated by AF9/ENL either through the reader function of YEATS domains or MLL fusions plays pivotal roles in carcinogenesis, suggesting a putative therapeutic window for AF9/ENL inhibitors in MLL-r leukemias.

A common rationale for the AF9/ENL inhibitor design exploits the tunnel-like property of the reader pocket and a unique π-π-π stacking interaction in YEATS proteins. The acetyllysine-binding pocket of YEATS domains shows an “end-open” characteristic, which is ideal to accommodate extended lysine acylation chains, such as *N*^*ε*^-propionyl-L-lysine (PrK, lysine propionylation), *N*^*ε*^*-*butyryl-L-lysine (BuK, lysine butyrylation), and *N*^*ε*^*-*crotonyl-L-lysine (CrK, lysine crotonylation) and the expanded reader activity allows YEATS domains to coordinate ligands with different structural moieties.^23-27^ Additionally, crystallography studies revealed AF9 and ENL bind preferentially to CrK over AcK via a π–π–π sandwich interaction with two highly conserved aromatic side chains F59 and Y78 and CH-π interaction with F28.^23-24, 28^ The enhanced binding affinity is believed to be attributed to the extra intermolecular force provided by the conjugated π system on the crotonyl group.^23-24^ These findings have led to the design and identification of a set of small molecule and peptide YEATS inhibitors.^27, 29-34^ However, as the residues involved in reader pockets of AF9 and ENL are identical, it is difficult for an inhibitor to differentiate between ENL and AF4 solely by targeting ligand-protein interactions. Despite their high sequence homology, the two close YEATS proteins AF9 and ENL appear to play different roles in cancers. For example, it was established that ENL, but not AF9 is required for acute leukemia progression.^4-5^ In addition, knockout studies showed the depletion of ENL had minimal effects on normal hematopoietic stem cells.^5^ These findings suggest that selective inhibition of ENL could be an effective and less toxic approach for leukemia therapeutics.

Compared to small-molecule drugs, peptides display improved affinity and selectivity owing to their ability to provide extra interactions with the surrounding area of active site. As showcased by JYX-3, this AF9 peptide inhibitor targets a proximal site outside the acyllysine-binding pocket, resulting in a remarkable selectivity for AF9 over ENL.^34^ Over the years, phage display has emerged as a robust tool to screen peptide inhibitors. However, traditional phage display is agnostic to where on the target protein the displayed peptide is bound, which often leads to the enrichment of nonproductive ligands. In order to identify selective ENL YEATS inhibitors in a high-throughput manner, we envisioned that the previously developed phage-assisted, active site-directed ligand evolution (PADLE) technique could be applied. PADLE relies on a genetically encoded noncanonical amino acid (ncAA) that binds the active site of a protein target to direct phage-displayed peptides for selection.^35^ To conduct PADLE on ENL YEATS, a warhead with known target-ligand interactions for ENL YEATS needs to be genetically incorporated into a phage-displayed sequence-randomized peptide library (Figure 1). This enables a rationally designed interaction to guide the displayed-peptides towards the active site, increasing the probability that a given selection will produce a productive inhibitor. In addition to exploiting a known binding interaction, the other randomized residues provide additional interactions to the areas adjacent to the binding pocket, increasing both affinity and selectivity for enriched ENL YEATS inhibitors. In the current reported work, we wish to report our progress in using PADLE in combination with the structure-activity relationship (SAR) study to identify a selective and cellularly potent peptidyl inhibitor for ENL YEATS.

**Figure 1.**
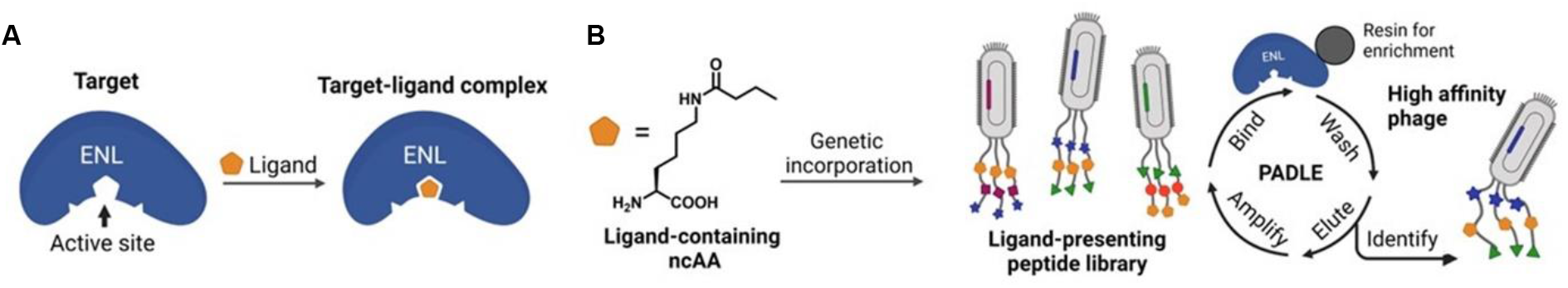
A schematic diagram of phage-assisted, active site-directed ligand evolution (PADLE) of ENL YEATS inhibitors. (A) A diagram that illustrates the interaction between ENL YEATS and a ligand. (B) The genetic incorporation of a ligand-fused ncAA into phage-displayed peptides for active site-directed binding to ENL YEATS that is followed by stringent wash and elution to select high-affinity and selective phages. Figures created with BioRender.com.

## Results and Discussion

### The genetic incorporation of butyryllysine (BuK) into phage-displayed peptides

For the identification of an ENL YEATS inhibitor through PADLE, a genetically incorporated warhead with known capabilities to bind the target protein is required. Ideally, CrK could serve as a suitable warhead as it can be recognized by YEATS domains. However, in previous research, we and others discovered that when an electrophilic ncAA (e.g., a Michael acceptor) is presented along with a cysteine in the phage displayed peptide, it can undergo proximity-driven cyclization.^36-37^ Since there is no stereochemical preference for this thiol-ene addition, the cysteine may engage the planar crotonamide group from both sides, resulting in the formation of two stereoisomers, adding complexity to ligand identification. Besides CrK, due to the end-open feature of reader pocket, YEATS domains can recognize several other histone acylation marks. It was reported that AF9 YEATS binds BuK with enhanced affinity compared to AcK.^23-24^ Although lacking a π system, the extended acyl chain provides extra binding force through a CH-π interaction with F28.^23-24^ Consequently, BuK was chosen as an alternative ncAA in our study. The incorporation of BuK in a phage displayed peptide has been described.^35^ Briefly, BuKRS, a BuK-tRNA synthetase evolved from MmPylRS, was used to produce a phage in which the gIII gene containedan amber codon at its *N*-terminal coding region. Titer results showed a nearly 100-fold difference in phage production in response to the absence and presence of BuK, demonstrating the successful incorporation of this ncAA into phage-displayed peptides (Figure S1).^35^

### The construction of a BuK-containing phage display library and the biopanning against ENL YEATS

To apply the PADLE-based ligand search for ENL, the BuK-presenting peptides (7-mer) were displayed on phage using the method previously developed.^35^ In short, the phages were produced in the presence of 5mM BuK using TOP10 cells harboring the following three plasmids: (1) pADL-gIII, which contains the 7-mer amber-obligate peptide-coding DNA library positioned at the *N*-terminus of M13 pIII gene, (2) pCDF-BuKRS, which encodes BuKRS and its cognate tRNA^Pyl^ for the genetic incorporation of BuK at amber codons in the 7-mer library, and (3) M13KO7TAA, a helper phage plasmid engineered to have a TAA mutation at pIII so that its lack of a functional pIII allows an efficient multivalent display of the 7-mer library. The ENL YEATS domain was recombinantly expressed in *E. coli* with an *N*-terminal AviSUMO-Tag in the presence of BirA, a biotin ligase, for site-specific biotinylation at the AviTag. The biotinylated AviSUMO-ENL was immobilized on streptavidin-coated magnetic beads for panning with the BuK-containing phages. The selection pressure was gradually increased by lowering the protein usage and increasing the number of washes to promote enrichment of potent binders. Through three rounds of panning, an approximately 362-fold increase in the total number of eluted phages was achieved in spite of these increased selection pressures, suggesting a successful selection. Phage libraries collected from the first and third rounds of selection were sequenced using Illumina next generation sequencing (NGS). Several R scripts were written and used to translate the DNA reads and filter off low-quality sequences using paired-end processing.^35^ The deep sequencing results from 95,763 reads showed that 7,124 unique sequences were obtained from the first round of selection (Figure 2A). Among all the sequences, only 102 of them contained more than 100 copies (blue area), while the remaining sequences all had low abundances (<10^2^ copies, dark and light gray areas), which constituted 83.8% of first round population. NGS data revealed that after three panning rounds, the selection converged to two highly enriched sequences: YDVYCYX (**ENL-S1**) and WWIIEXG (**ENL-S2**, X denotes BuK). Out of 1,346 different unique sequences collected from the third round, **ENL-S1** and **ENL-S2** represented 82.5% of the entire library (Figure 2A and Figure S5-6).

**Figure 2.**
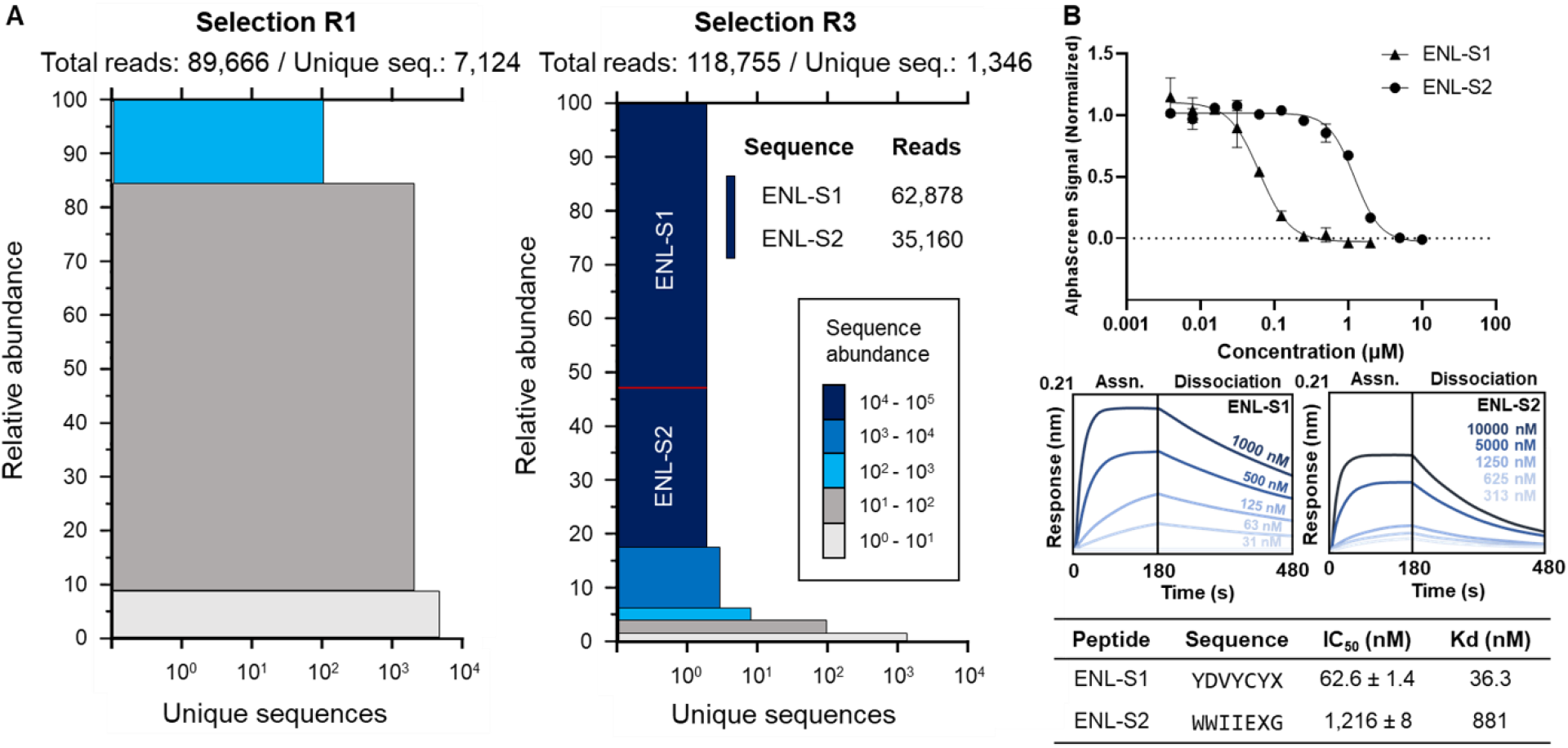
Phage panning identified two peptide ligands for ENL YEATS. (A) The visualization of next-generation sequencing results from first and third panning rounds. Each colored block represents all sequences with the specific abundance. The block height corresponds to the percent abundance of sequences in each sector, and the width represents the number of unique sequences per sector. After three rounds of panning, the library converged into two highly enriched sequences, **ENLS1** (YDVYCYX) and **ENLS2** (WWIIEXG). X denotes *N*^*ε*^-butyryl-L-lysine. (B) Binding and inhibition parameters of the selected peptides. Binding kinetics are characterized with Biolayer Interferometry with varying concentrations of selected peptides. The nonlinear least squares-fitting curves are represented. IC_50_ values are measured with AlphaScreen assays and given as the mean ± standard deviation (SD) of three individual experiments (n = 3).

### Development of selective ENL YEATS inhibitors

To test the inhibitory potency of the two selected peptides for the ENL YEATS domain, **ENL-S1** and **ENL-S2** were chemically synthesized through solid-phase peptide synthesis (SPPS) and evaluated by an AlphaScreen assay. In this assay, a biotinylated H3K27cr peptide and a 6x His-tagged ENL YEATS domain (His-ENL) were immobilized on PerkinElmer streptavidin donor and nickel-chelating (Ni-NTA) acceptor beads, respectively. The use of a His-ENL instead of the original AviSUMO-ENL construct in the assay prevents potential false positive results derived from ligands which bind to the SUMO protein but not to the ENL YEATS domain. Results in Figure 2B demonstrated that both **ENL-S1** and **ENL-S2** inhibited interactions between ENL YEATS and H3K27cr with an IC_50_ value as 63 and 1,216 nM, respectively. We then assessed the binding affinity of two ligands for ENL YEATS using biolayer interferometry (BLI). Briefly, His-ENL was immobilized onto Ni-NTA functionalized biosensors to measure the binding kinetics of target peptides at different concentrations. Results showed **ENL-S1** bound strongly to ENL with a determined dissociation constant *K*_*d*_ of 36.3 nM, whereas **ENL-S2** bound severalfold weaker to the target protein with a *K*_*d*_ value of 881 nM (Figure 2B and Figure S6). For **ENL-S1** and **ENL-S2**, their determined *k*_*on*_ and *k*_*off*_ values were 6.23 × 10^4^, 6.29 × 10^3^ M^-1^ s^-1^ and 2.26 × 10^−3^, 5.54 × 10^−3^ s^-1^, respectively. Compared to **ENL-S2, ENL-S1** has faster binding to ENL YEATS and slower release from it, indicating its potential long term action on the ENL YEATS inhibition. Collectively, these data confirmed that **ENL-S1** displayed much better potency than **ENL-S2**, making it an ideal candidate for further optimization as a high-affinity ENL inhibitor.

AF9 and ENL YEATS domains bind preferentially to CrK over AcK via a π–π–π sandwich interaction with two aromatic residues F59 and Y78 in YEATS (positions are corresponding to that in ENL YEATS).^23-24^ Since the π–π interactions are not available to butyryl group due to its lack of a double bond, we envisioned that replacing the BuK residue in **ENL-S1** with a conjugated π system will lead to a more potent inhibitor through the enhanced stacking interactions. In previous research, two modified H3 peptides with *N*^ε^*-*2-furancarnonyl-L-lysine (FurK) and *N*^ε^*-*5-oxazolecarbonyl-L-lysine (OxaK) that contained expanded π systems to provide extra intermolecular interactions with the AF9 YEATS domain showed a 4.5- and 14.7-fold binding enhancement compared to that with CrK.^33^ Inspired by this finding, we synthesized three **ENL-S1** derivatives: **ENL-S1c, ENL-S1f**, and **ENL-S1o** with crotonyl, 2-furancarbonyl, and 5-oxazolecarbonyl replacing the butyryl group in **ENL-S1** (Figure 3C). As predicted, all three derivatives **ENL-S1c, ENL-S1f**, and **ENL-S1o** showed increased potency compared to **ENL-S1** with IC_50_ values as 20, 22, and 10 nM, respectively (Figure 3A), supporting the critical role of a conjugated π system in achieving strong binding to a YEATS domain. Notably, the correlation between substituents and binding affinity also suggests that BuK in **ENL-S1** is positioned in the π–π–π sandwich cage of ENL YEATS to guide displayed peptides to the active site. All three inhibitors were also characterized using BLI. They all have determined *K*_*d*_ values below 20 nM (Figure S7). Among them, **ENL-S1o** has a remarkable low *K*_*d*_ value of 2.0 nM. As far as we know, this is the strongest inhibitor that has been developed for the ENL YEATS domain. All three inhibitors displayed fast binding to ENL YEATS and slow release from it, indicating their potential long-term action for ENL YEATS inhibition.

**Figure 3.**
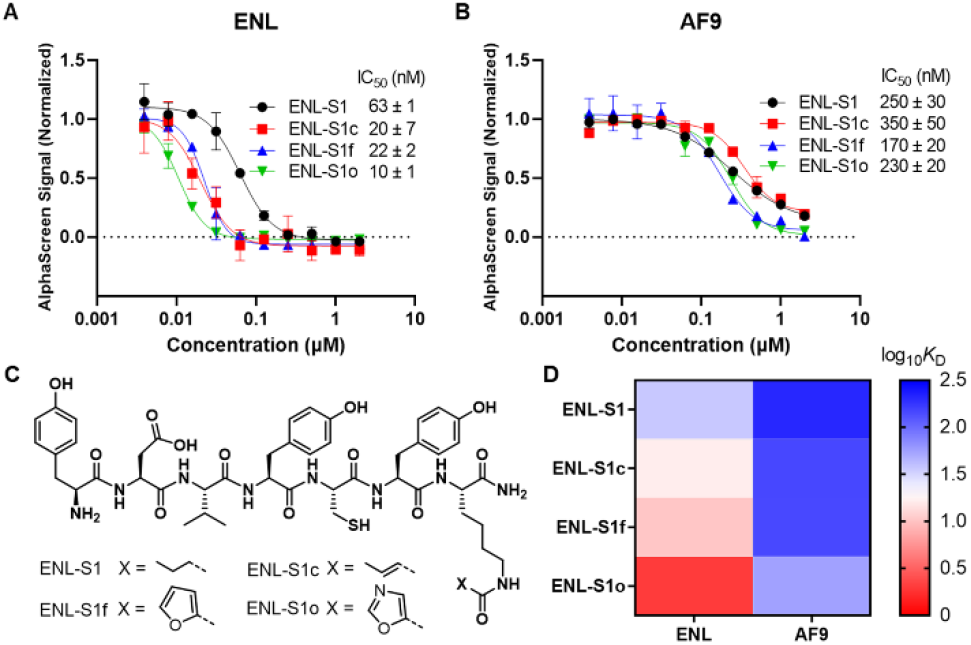
Characterization and optimization of **ENL-S1**. (A,B) AlphaScreen analysis of IC_50_ of **ENL-S1** and its derivatives in inhibition of (A) ENL and (B) AF9 YEATS binding to the corresponding acylated histone peptide. (C) Chemical structures of **ENL-S1** and three derivatives developed for targeting the π-π-π stacking for YEATS inhibition. (D) BLI measurement for the binding affinities of **ENL-S1** and its derivatives with AF9 and ENL YEATS. IC_50_ values are given as the mean ± SD, n = 3.

These ultrapotent inhibitors were then tested to confirm their selectivity for ENL over other YEATS proteins. Selective targeting ENL is preferred in leukemia interventions as it plays a pivotal role in the progression and maintenance of several AMLs and poses minimal harm on normal hematopoietic stem cells upon inhibition. The inhibition assays performed with AF9 YEATS domain showed that **ENL-S1** and all three derivatives are more selective toward ENL over AF9 YEATS (Figure 3B). Judging from IC_50_ values, **ENL-S1, ENL-S1c, ENL-S1f**, and **ENL-S1o** displayed 4.0-, 18-, 7.7-, and 23-fold lower potencies toward AF9 YEATS compared to ENL YEATS. These results show that despite the high sequence and structure similarity between AF9 and ENL YEATS domains, our identified peptides are able to distinguish between these two highly conserved domains. We believe the specificity arises from the peptide residues engaging surface areas adjacent to the active site of ENL YEATS to provide selective interactions. This showcases the practicality of the PADLE-based approach to develop potent and selective inhibitors. Similarly, BLI analyses were conducted to reaffirm preferential binding of inhibitors to ENL YEATS. In agreement with the results from inhibition assay, all four peptides showed selectivity toward ENL over AF9 (Figure 3D and Figure S7). In particular, **ENL-S1** (*K*_*d*,ENL_ = 36.3 nM, *K*_*d*,AF9_ = 205 nM) and **ENL-S1c** (*K*_*d*,ENL_ = 16.2 nM, *K*_*d*,AF9_ = 147 nM) displayed 5.6- and 9.1-fold higher affinities for ENL YEATS. Two peptides with extended π systems, **ENL-S1f** (*K*_*d*,ENL_ = 10.3 nM, *K*_*d*,AF9_ = 144 nM) and **ENL-S1o** (*K*_*d*,ENL_ = 2.0 nM, *K*_*d*,AF9_ = 55.1 nM) showed even higher, preference for targeting ENL YEATS, as they bound 14.0- and 27.6-fold, respectively, stronger affinity toward ENL than AF9. Overall, these data demonstrate our peptides preferentially target ENL over AF9.

### A SAR study to search for inhibitors with potential cellular permeability

Although **ENL-S1** and its derivatives showed promising results in *in vitro* assays, their 7-mer peptide nature leads to a concern of their access to the nucleus, where ENL presents. Therefore, to improve the physicochemical properties of **ENL-S1**, an alanine scan was performed to study its SAR (Figure 4 and Figure S8). The analysis showed that replacing the 2^nd^ residue with alanine slightly improved inhibition. Replacing the two tyrosine residues at the 1^st^ and 4^th^ positions resulted in similar decrease in inhibition. Mutation of the 3^rd^ and 5^th^ residues caused dramatic decrease in activity, suggesting these residues were crucial for ENL binding. As predicted, replacing BuK with alanine abolished the inhibition, confirming its vital role in the molecular recognition. Interestingly, substituting the 6^th^ residue also resulted in complete loss of activity, indicating this tyrosine is critical for the binding to ENL YEATS. According to the alanine scan results where the displacement of aspartic acid at the second position exhibited little perturbation to inhibition and only a moderate decrease was observed when the first tyrosine was replaced, we removed the first two amino acids while retaining the rest essential residues. In addition, an acetyl protecting group was added to the *N*-terminus to prevent N-terminal degradation and to increase peptide permeability. Subsequently, four inhibitors, **tENL-S1, tENL-S1c, tENL-S1f**, and **tENL-S1o** were generated for further characterizations (Figure 5A).

**Figure 4.**
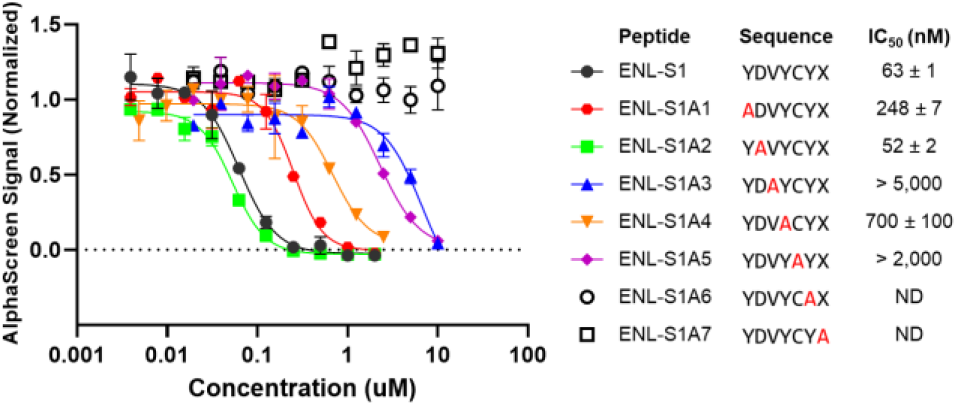
Structure-activity relationship studies of **ENL-S1** to identify key residues. An alanine scan was performed by iteratively replacing each residue with an alanine and testing for change of ENL inhibition through AlphaScreen assays. IC_50_ values are given as the mean ± SD, n = 3.

**Figure 5.**
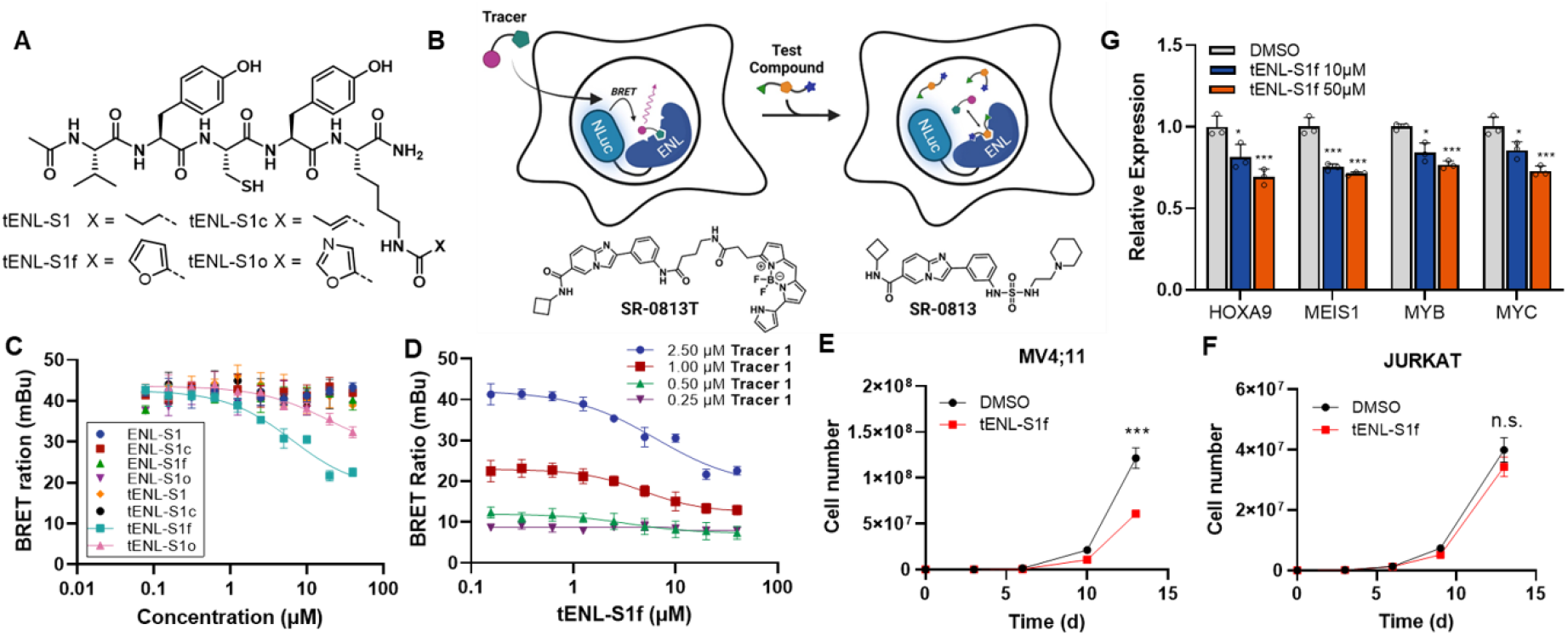
(A-D) Development of a NanoBRET assay for cellular target engagement profiling of ENL inhibitors. (A) Chemical structures of truncated **ENL-S1** derivatives. (B) Schematic of the NanoBRET assay for ENL YEATS. A NanoLuc luciferase (NLuc)-ENL YEATS fusion protein is expressed in HEK293T cells. **SR-0813T** (bottom left) developed from an ENL inhibitor SR-0813 (bottom right) is used to achieve BRET. Upon **SR-0813T** binding to ENL YEATS, luminescence generated from NLuc is transferred to the tracer fluorophore, resulting in a fluorescence emission. The test compounds compete with **SR-0813T** for binding to ENL YEATS and leads to an attenuated fluorescence signal. (C) NanoBRET curves of **ENL-S1** and its derivatives displaying their relative affinities for ENL YEATS in HEK293T cells. (D) Measurement of apparent intracellular affinity of **tENL-S1f** for ENL-YEATS. (E-G) On-target effects of **tENL-S1f** in leukemia. Cell growth inhibition of **tENL-S1f** in (E) ENL-dependent MLL-fusion leukemia cell line and (F) ENL-insensitive acute leukemia cell line. Mean ± SD, n = 3, P values by two-tailed Student’s t-test to DMSO sample. (G) qPCR analysis of transcript abundance in MOLM13 cells treated with **tENL-S1f** (normalized to DMSO). Raw BRET ratios in (C,D) are shown as mean ± SD, n = 3. Data in (E-G) represent Mean ± SD, n = 3, P values by two-tailed Student’s t-test to DMSO sample. *P < 0.05, **P < 0.01, ***P < 0.001. Not significant (n.s.) P > 0.05. Schemitic of NanoBRET design was created with BioRender.com.

### Validation of ENL inhibitors by NanoBRET assays

With the truncated peptides in hand, we then evaluated whether these inhibitors could engage ENL YEATS in cells using a NanoBRET assay.^38^ This technique, which measures the competitive binding between a fluorescent tracer and test compound through a bioluminescence resonance energy transfer (BRET), has emerged as a robust and sensitive tool to assess ligand engagement in cellular environments. There are three components required to gauge the peptide inhibitory potency for ENL through NanoBRET assays: (1) an ENL YEATS-NanoLuc luciferase fusion protein, (2) a NanoLuc luciferase substrate, and (3) a fluorescent tracer that binds specifically and reversibly to the active site of ENL YEATS. During the assay, the substrate furimazine reacts with NanoLuc luciferase to produce bioluminescence, which is quenched by the nearby tracer bound to ENL-NanoLuc fusion via BRET. The affinity of peptides can then be measured through the decrease in BRET as test compounds engaging ENL and displacing the tracer. Previously, a NanoBRET assay based on Histone 3.3 (H3.3) was developed for YEATS proteins.^29^ In which, the NanoLuc luciferase was fused with a full-length AF9 or ENL and co-expressed with histone H3.3-HaloTag. The histone H3.3 was expected to be acetylated through post-translational modifications and recognized by NanoLuc-AF9/ENL. The NanoLuc-AF9/ENL:H3.3 complex was treated with a HaloTag NanoBRET tracer ligand to achieve BRET. While it was successfully used to confirm the cellular activity of YEATS inhibitors against AF9, no responses were observed for ENL upon inhibitor treatment. This was likely attributed to the difference in binding affinity between AF9 and ENL to histone acetylation marks; as shown in isothermal titration calorimetry (ITC) analyses, the ENL YEATS domain binds overall weaker to the acetylated H3 peptides than its paralog AF9 YEATS (ENL: *K*_*d*, H3K9ac_ = 32.2 μM, *K*_*d*, H3K18ac_ = 50.0 μM, *K*_*d*, H3K9_ = 30.5 μM; AF9: *K*_*d*, H3K9ac_ = 3.7 μM, *K*_*d*, H3K18ac_ = 11.0 μM, *K*_*d*, H3K27ac_ = 7.0 μM).^1, 4^ Since BRET signal is dependent on the ligand-protein association, having an inadequate interaction between histone H3.3 and a YEATS domain may result in a small assay window and poor applicability.

In light of this, we developed a NanoBRET assay based on **SR-0813T**,^39^ a small-molecule tracer adapted from a highly potent ENL inhibitor SR-0813 (Figure 5B).^31^ To achieve BRET, a fluorophore capable of accepting luminescent energy generated from NanoLuc is needed. An ideal choice is the boron-dipyrromethene (BODIPY) dyes owing to their excellent photophysical properties, such as high fluorescence/absorption levels and relative insensitivity to the changes in solvent pH or polarity.^40^ The lack of charges and molecular polarity on BODIPY structure also favors its usage in tracer design as it minimizes the chance of disrupting functional properties of tagged ligand upon installation.^41^ As such, a pyrrolyl substituted BODIPY (Py-BODIPY) was appended to the SR-0813 scaffold via a short linker, 4-aminobutyric acid, to afford **SR-0813T**. The use of Py-BODIPY provided optimal spectral separation between donor and acceptor emissions with its red-shifted emission wavelength. Meanwhile, we also established a stable cell line for NanoLuc-ENL expression. In short, a NanoLuc-ENL fusion was constructed by cloning cDNA encoding ENL YEATS domain (aa 1-148) into a NanoLuc containing vector pCDH-EF1-Nluc. The plasmid was transduced into HEK293T cells using lentiviruses. A selection with increasing amount of puromycin over two weeks was carried out to afford the final stable cells. To validate the binding of **SR-0813T** to the YEATS domain we performed a fluorescence polarization assay with ENL YEATS (Figure S9). **SR-0813T**, compared to H3 peptides, exhibited an improved binding affinity for ENL YEATS (*K*_*d*_ = 1.1 μM), albeit weaker than its parental compound SR-0813, presumably due to the chemical modifications introduced by conjugation.

Using the developed ENL-NanoBRET assay, we characterized cellular potency of our developed peptidyl inhibitors of ENL YEATS. The analysis was done with 2.5 μM of **SR-0813T** and varying concentrations of inhibitors (Figure 5C). All linear 7-mer peptides showed no detectable inhibition against ENL YEATS within the assay window. Two of the four truncated **ENL-S1** derivatives exhibited ENL engagement; **tENL-S1f** showed a clear dose-dependent displacement of **SR-0813T** from NanoLuc-ENL (IC_50_ = 6.4 ± 1.6 μM), whereas **tENL-S1o** showed little cellular activity with ∼50% inhibition at 40 μM. The apparent intracellular affinity of **tENL-S1f** was further evaluated by NanoBRET at various tracer concentrations (Figure 5D). Overall, the peptide inhibited ENL with IC_50_ values in the low-micromolar range in cells (IC_50, 0.5μM_ = 4.1 ± 1.1 μM, IC_50, 1.0μM_ = 4.2 ± 1.1 μM, IC_50, 2.5μM_ = 6.4 ± 1.6 μM). BLI analysis of **tENL-S1f** confirmed that the inhibitor is selective toward ENL YEATS over three other human YEATS proteins with a 12.0-fold selectivity over AF9 and no binding to YEATS2 and GAS41 up to 10 μM **tENL-S1f** used (Figure S10). These results suggested that the truncation of the 7-mer peptide **ENL-S1f** to afford **tENL-S1f** allowed for a better cellular target engagement while retaining the selectivity of resultant peptide, although at the cost of its potency.

### tENL-S1f inhibits leukemia cell growth and downregulates the expression of ENL target genes

Encouraged by the cellular studies, we next examined the effects of **tENL-S1f** on acute leukemia proliferation in an MLL-rearranged cell line MV4;11 (MLL-AF4 AML) whose growth is sensitive to ENL depletion.^5^ We found that **tENL-S1f** exhibited ∼50% inhibition of MV4;11 cell growth at 10 μM after 14 days of treatment (Figure 5E). In contrast, the growth pattern of JURKAT cells (T-cell ALL), an ENL-independent cell line,^5^ showed no detectable response to the treatment with **tENL-S1f** (Figure 5F). As aforementioned, ENL drives leukemogenesis through the ENL YEATS-chromatin association at oncogenic gene loci. To confirm whether our peptide could disrupt the ENL YEATS-chromatin interaction and thereby downregulates oncogene expression. We evaluated the relative mRNA level of several ENL-target genes, including *HOXA9, MEIS1, MYB* and *MYC* in MOLM-13 cells (MLL-AF9 AML) treated with **tENL-S1f**. Quantitative PCR results showed all genes were suppressed in response to **tENL-S1f** treatment at 10 μM. At 50 μM, a ∼30% loss in transcription levels was observed for all four leukemic genes (Figure 5G). Altogether, these results indicate **tENL-S1f** exhibits on-target effects, suppresses ENL target gene expression, and ultimately inhibits ENL-dependent leukemia proliferation.

### MD simulations to predict the interactions between tENL-S1f and ENL YEATS

To further understand how tENL-S1f engage YEATS proteins, we performed molecular dynamic (MD) simulations using structures of AF9 YEATS bound to H3K9Ac peptide (PDB 4TMP) and ENL YEATS bound to H3K27Ac peptide (PDB 5J9S) as templates. In AF9 simulation, the predicted structure showed **tENL-S1f** bound to the substrate-binding site of AF9 YEATS (Figure S11). Particularly, F59 involves a π-π interaction with the furanyl group in **tENL-S1f**. Three residues S58, Y78, and A79 interact with FurK side chain. G80 located at the end of reader pocket and L108 at the lower end of the substrate recognition region form several backbone hydrogen bonds with **tENL-S1f**. In comparison, ENL simulation results showed tENL-S1f engages the active site of ENL with a different conformation from the binding to AF9 (Figure 6A). The key interactions are highlighted in Figure 6B. Two aromatic residues F59 and Y78 formed π-π stacking interactions with the furan ring in **tENL-S1f**. G80, L106, and L108 formed several hydrogen bonds with the **tENL-S1f** backbone. Additionally, residue H30 forms an extra π-stacking with Y2 of **tENL-S1f**, and D103 forms a hydrogen bond with Y4 of **tENL-S1f** – neither interaction was found in AF9 simulation. Collectively, the simulation results provided a possible explanation for the high affinity of **tENL-S1f** and its selectivity for ENL YEATS over AF9 YEATS.

**Figure 6.**
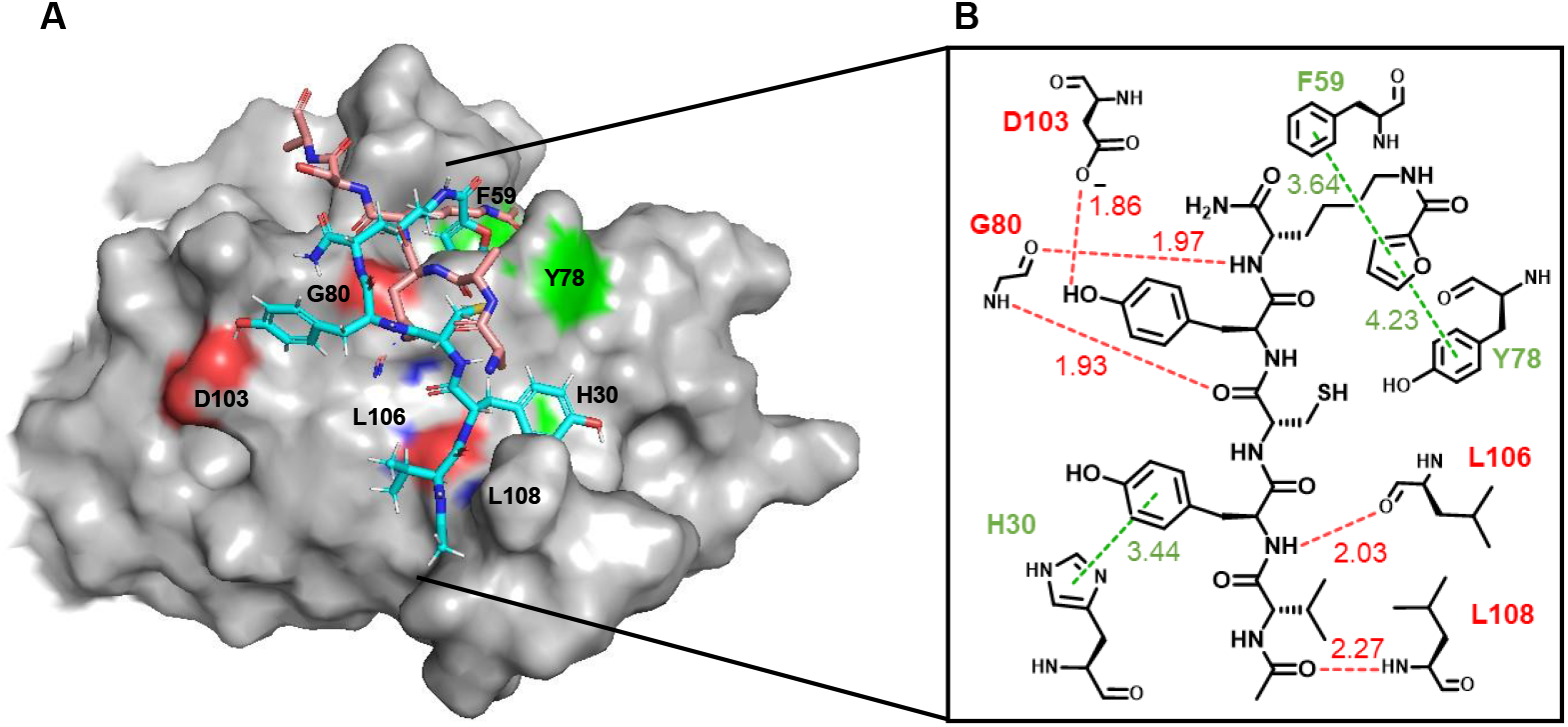
Molecular dynamic simulations to predict the interactions between **tENL-S1** and ENL YEATS. (A) The MD simulation predicted complex structure of **tENL-S1** (cyan) bound to the substrate binding site of ENL YEATS (gray). The native substrate H3K27Ac peptide is shown in salmon. (B) Interactions between **tENL-S1f** and ENL YEATS predicted by molecular dynamic simulations. π-stacking (green) and hydrogen-bond (red) interactions are shown in dash lines with distance indicated in angstrom. MD simulations were performed based on the reported crystal structure (5J9S).

## Conclusion

Epigenetic proteins are a rich source of potential therapeutic targets. Recently, a set of epigenetic drugs, e.g. tazemetosta and vorinostat, targeting epigenetic writers and erasers including histone methylases and histone deacetylases have been developed and applied in clinical use.^42-43^ Unlike writers and erasers which utilize a distinct binding site to catalyze post-translational modifications, reader proteins typically provide a cavity or binding groove with a large surface area to engage chromatin, rendering them difficult to be targeted by small-molecule drugs.^44^

Our PADLE technique has proven to be viable in generating potent and selective peptidyl inhibitors through the genetic incorporation of a known ligand into a peptide library. In this study, we successfully applied BuK as the warhead to direct the displayed peptides towards the active site of ENL YEATS and identified a peptide inhibitor **ENL-S1**. The butyryl group in **ENL-S1** was substituted with moieties carrying a conjugated system to exploit the π-π-π stacking interactions, which resulted in enhancements in both inhibitory potency and binding affinity, supporting the hypothesis that BuK is positioned in the π-π-π sandwich cage and used to guide peptides toward the active site. Notably, due to the selective interactions formed between the peptide residues and the surrounding area of reader pocket, **ENL-S1** and its derivatives exhibited remarkable selectivity for ENL YEATS over AF9 YEATS. These peptides were optimized based on alanine scan results to acquire better cell permeability. The resultant peptides were evaluated by a developed NanoBRET assay for their cellular target engagement, in which a low-micromolar ENL inhibitor **tENL-S1f** was identified. Further analyses confirmed that **tENL-S1f** is selective for ENL over other YEATS domains and exhibits on-target effects in inhibiting ENL target gene expression in cells and leukemia cell growth.

As the first of its kind study, the current work demonstrated the viability of using PADLE for the development of peptidyl inhibitors for an epigenetic reader. Over the past decades, through the use of genetic code expansion, we and others have successfully incorporated a wide variety of ncAAs, including acylated and methylated lysines.^45-49^ As these ncAAs often serve as histone marks and provide well-defined target-ligand interactions with epigenetic proteins (e.g., histone deacetylases, bromodomains, and chromodomains), such interactions can be well exploited in a high-throughput manner via PADLE to facilitate the identification of potent and selective peptide therapeutics. We believe the PADLE technique with its general applicability will find broad applications in this aspect.

