## Supporting Information for "Phage-Assisted, Active Site-Directed Ligand Evolution with a Genetically Encoded *N*^*ε*^-Butyryl-L-Lysine to Identify a Cellularly Potent and Selective Inhibitor for the ENL YEATS Domain as an Anti-Leukemia Agent"

### Supplementary Methods

#### Chemical Synthesis

<u>*The Synthesis of BuK*</u>:

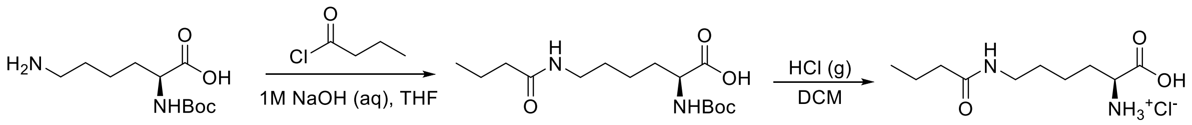


Scheme S1. Synthesis of N$^{\varepsilon}$-butyryl-L-lysine hydrochloride (BuK).

The synthetic route shown in Figure S1 was adapted from Gattner et al.[1]
Boc-Lys-OH (15 g, 61 mmol) was dissolved in 1M NaOH (150 mL) and THF (150 mL) and cooled on ice. Butyryl chloride (6.3 mL, 62 mmol) was added dropwise and stirred overnight at room temperature. To prevent polymerization, the reaction was concentrated under reduced pressure at 0 °C for the removal of THF. The aqueous solution was washed with methyl tert-butyl ether (3 x 75 mL), and then the aqueous layer was acidified using 6 N HCl. The aqueous solution was extracted with EtOAc (3 x 75 mL), and the combined organic layers were washed with brine (75 mL), dried over $Na_2SO_4$, filtered, and concentrated under vacuum to give Boc-BuK-OH (18 g, 94% yield), which was used immediately in the next step. Boc-BuK-OH (18 g, 57.0 mmol) was dissolved in DCM (100 mL), and attached via cannula to a separate 3-neck flask charged with NaCl (45 g, 770 mmol). Sulfuric acid (40 mL, 750 mmol) was added to the NaCl dropwise via addition funnel. The resulting HCl was slowly bubbled into the solution of Boc-BuK-OH and stirred until bubbling ceased. The resulting product was dissolved in water (100 mL), leaving behind insoluble polymer that formed. The aqueous layer was flash frozen and lyophilized to give $N^{\varepsilon}$-butyryl-L-lysine hydrochloride (BuK-HCl, 9.3 g, 65% yield) as a white solid. $^1$H NMR ($D_2O$) $\delta$ = 4.03 (t, J = 6.30, 1H), 3.18 (t, J = 6.82, 2H), 2.18 (t, J = 6.72, 2H), 1.85-2.02 (m, 2H), 1.33-1.62 (m, 6H), 0.87 (t, 3H, J = 7.41).

<u>*Synthesis of Fmoc-Protected Lysine Derivatives*</u>:

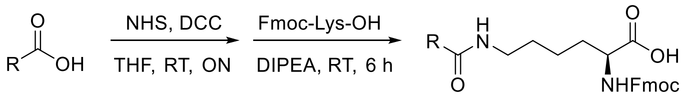


Scheme S2. Synthesis of Fmoc-protected lysine derivatives

The synthetic route shown in Figure 4.6 was adapted from Li et al.[2]
Carboxylic acid (1.5 eq.) and NHS (1.4 eq.) were dissolved in dry THF. DCC (1.4 eq.) in dry THF was added into the above solution and stirred at room temperature overnight. The reaction mixture was filtered and to the filtrate was added Fmoc-Lys-OH (1 eq.) together with DIEA (3 eq.). The resulting reaction mixture was allowed to stir at room temperature for another 6 h. The pH of the mixture was adjusted to 7 with 1 M HCl. The solvent was *in vacuo*. The residue was extracted with DCM and 1 M HCl. The organic layer was washed by brine and dried over $Na_2SO_4$. After removal of solvent, the crude product was purified by silica gel column chromatography.

**Solid Phase Peptide Synthesis**

Peptides were synthesized on ProTide Rink amide low loading resin (CEM #R002) using an automated parallel peptide synthesizer MultiPep 2. Noncanonical amino acids were installed using Fmoc-protected lysine derivatives (5 equivalents). Fmoc-amino acids were deprotected using 20% piperidine in DMF. Free Fmoc-amino acids (5 equivalents) were then coupled with 4 M *N*-methylmorpholine fand 0.5 M HATU in DMF for two 30-minute cycles. The *N*-terminal acetylation was done with 3 mL of 25% acetic anhydride in DCM for 5 min at room temperature. Peptides were cleaved from resin by agitating for 3 hours with 2 mL of cleavage solution (92.5:2.5:2.5:2.5 TFA:$H_2O$:DODT:TIS). The cleavage solution was filtered and precipitated into 40 mL of cold ether, and the precipitated peptides were pelleted (4,000 rcf, 10 min, 4 °C), the supernatant decanted, and the pellets were dissolved in DMF and purified via reverse-phase semi-preparative HPLC (acetonitrile in water with 1% formic acid, 18 mL/min, PDA) using a Shimadzu Shim-pack GIS C18 column 10 m (25mm x 250 mm, Shimadzu #227-30115-04).

**Protein Expression and Purification**

<u>*Expression of Avi-SUMO*</u>:
A stop codon TGA was introduced to the *C*-terminus of SUMO protein in pET28a-AviSUMO plasmid (provided by Dr. Pingwei Li at Texas A&M University). The plasmid was co-transformed with pBirAcm (provided by Dr. Pingwei Li at Texas A&M University) into BL21(DE3pLysS) chemically competent cells. These cells were grown in 1 L of 2xYT medium with chloramphenicol (34 μg/mL) and kanamycin (50 μg/mL) until $OD_{600}$ = 0.5-0.8, at which point the expression was induced with 0.4 mM IPTG in the presence of 50 μM biotin at 16 °C for 18 hours. Cells were pelleted (4000 rcf, 10 minutes) and resuspended in 50 mL of lysis buffer (50 mM Tris, 300 mM NaCl, 10 mM imidazole, pH 7.8) with 0.1 mM phenylmethylsulfonyl fluoride (PMSF). Cells were lysed using sonication, cellular debris was pelleted (16,000 rcf, 30 minutes, 4 °C), and the lysate was transferred to 2.0 mL of Ni-NTA resins and incubated at 4 °C for 45 minutes. The protein-resin slurry was washed three times with wash buffer 1 (50 mM Tris, 300 mM NaCl, 30 mM imidazole, pH 7.8), three times with wash buffer 2 (50 mM Tris, 300 mM NaCl, 60 mM imidazole, pH 7.8), eluted with 6.0 mL of elution buffer (50 mM Tris, 300 mM NaCl, 250 mM imidazole, pH 7.8), dialyzed into storage buffer (50 mM Tris, 300 mM NaCl, 5 mM DTT, pH 7.8), and stored in aliquots at -80 °C.

<u>*Expression of Avi-SUMO-ENL*</u>:
The complementary DNA (cDNA) encoding ENL YEATS (aa 1−148) was cloned in pET28a-AviSUMO expression vector. The expression and purification of Avi-SUMO-ENL followed the same procedure as above for all proceeding steps.

<u>*Expression of His-ENL*</u>:
The cDNA encoding ENL YEATS (aa 1−148) was cloned in pET19b expression vector. The plasmid was transformed into BL21(DE3pLysS) chemically competent cells. These cells were grown in 1 L of 2xYT medium with ampicillin (100 μg/mL) until $OD_{600}$ = 1.0, at which point the expression was induced with 0.4 mM IPTG at 16 °C for 18 hours. The purification of His-ENL followed the same procedure as above for all proceeding steps.

#### Preparation and Expression of the Phage Display Library

*Expression and Purification of BuK-Incorporated M13 Phages*:

The affinity selection against ENL YEATS protein was performed using a TAG-enriched pADL-$(NNK)_7$ library developed by previous group members. Briefly, an $(NNK)_7$ library was introduced into the phagemid pADL-10b acquired from Antibody Design Labs at the *N*-terminus of pIII, and in order to enrich the randomized library for TAG-containing clones, a two-step superinfection-immunity-based selection was performed.[3]

The pADL$(NNK)_7$ library was transformed into electrocompetent TOP10 *E. coli* cells containing pEVOLCloDF-PylT-BuKRS (pEVOL-PylT-BuKRS with a CloDF origin of replication) and M13KO7(pIII-) (M13KO7 helper phage with a nonsense TAA mutation in gIII). Transformed cells ($6.2 \times 10^9$ transformants) were added to 1.1 L of fresh 2xYT with ampicillin (100 µg/mL), chloramphenicol (34 µg/mL), and kanamycin (25 µg/mL) and incubated at 37 °C until $OD_{600}$ = 0.5-0.8, at which point 1 mM IPTG, 5 mM nicotinamide, and 0.2% arabinose were added to induce phage expression. 100 mL of cells were added to a sterile flask to express the negative control, while 5 mM of BuK-HCl was added to the remaining 1 L of cells. Phages were expressed at 30 °C for 16 hours. Cells were pelleted via centrifugation (4000 rcf, 20 minutes, 4 °C), and the supernatant was poured into 5x precipitation buffer (2.5M NaCl, 20% PEG-8000). The phages were precipitated at 4 °C for two hours. Phages were pelleted via centrifugation (10000 rcf, 25 minutes, 4 °C), and the supernatant was discarded. Phages were resuspended in 30 mL of phage binding buffer (50 mM HEPES, 137 mM NaCl, 2.7 mM KCl, 1 mM $MgCl_2$, pH 8.0), centrifuged to clarify (4000 rcf, 20 minutes, 4 °C), 7.5 mL of 5x precipitation buffer was added, and phages were precipitated at 4 °C. Phages were once again pelleted via centrifugation (10000 rcf, 25 minutes, 4 °C), the supernatant decanted, and phages were dissolved in 1 mL of phage binding buffer. Phages were centrifuged to clarify (14000 rcf, 10 minutes, room temperature), and the phage solution was heat shocked at 65 °C for 15 minutes to kill any remaining cells. Phages were quantified through titering and stored at 4 °C until used for selection.

#### Affinity Selection Against ENL YEATS Protein

*Selection of BuK-Presenting Phages Against Avi-SUMO-ENL*:

50 µL of Sera-Mag streptavidin-coated magnetic beads (Cytiva) were washed three times with 1 mL of phage binding buffer (50 mM HEPES, 137 mM NaCl, 2.7 mM KCl, 1 mM $MgCl_2$, pH 8.0). 10 µg of AviSUMO-ENL in 1 mL of binding buffer was added to the beads and incubated at room temperature for 30 minutes under slow rocking. The supernatant was removed, and beads were washed three times with 1 mL of phage binding buffer and resuspended in 1 mL of blocking buffer (binding buffer with 1% BSA and 0.1% Tween-20. At the same time, 250 µL of 5x blocking buffer (binding buffer with 5% BSA and 0.5% Tween-20) was added to 1 mL of phage solution. Both mixtures were incubated at room temperature for 30 minutes under slow rocking. After 30 minutes, the blocking buffer was removed from the magnetic beads and the blocked phages were added, and the mixture was incubated for 30 minutes under slow rocking. Phages were then removed, and the beads were washed three times with 1 mL of wash buffer (blocking buffer with 0.1% Tween-20). The beads were transferred to a fresh tube after the first wash. 100 µL of elution buffer (50 mM glycine pH 2.2) were added to the beads, which were gently agitated for 15 minutes. The elution solution was removed from the beads and neutralized into 50 µL of neutralization buffer (1 M Tris pH 8.0). 25 mL of 2xYT with tetracycline (10 µg/mL) were inoculated with ER2738

cells and grown to $OD_{600}$ = 0.5-0.8. 5 mL of cell culture was removed from the cell culture and mixed with 10 µL of eluted phages to quantify total elution through titering. The remaining 20 mL of ER2738 cells were incubated with 140 µL of the phage solution for 45 minutes at 37 °C, and then centrifuged (4000 rcf, 10 minutes, 4 °C). The supernatant was discarded, and the cells were resuspended in 500 mL of 2xYT with tetracycline (10 µg/mL) and ampicillin (100 µg/mL) and grown overnight at 37 °C. Amplified cells were harvested and the phagemid library was extracted using a Miniprep kit (QIAGEN). The library was retransformed into electrocompetent Top10 *E. coli* cells containing pEVOL-BuKRS and M13KO7TAA to express phages for the next proceeding round.

For the 2nd and 3rd panning round, a negative control was added to the procedure. 100 µL of Sera-Mag streptavidin-coated magnetic beads were washed three times with 1 mL of phage binding buffer and split into two tubes. 10 µg of AviSUMO in 1 mL of binding buffer was added to one tube, and 10 µg of AviSUMO-ENL was added to the other (hereafter referred to as -ENL and +ENL, respectively). The bead/protein mixtures were incubated at room temperature for 30 minutes under slow rocking. The supernatants in both tubes were removed, and beads were washed three times with 1 mL of phage binding buffer and resuspended in 1 mL of blocking buffer. At the same time, 250 µL of 5x blocking buffer was added to 1 mL of phage solution. All three mixtures were incubated at room temperature for 30 minutes under slow rocking. After 30 minutes, the blocking buffer was removed from the -ENL tube and the blocked phages were added, and the mixture was incubated for another 30 minutes under slow rocking. Then the blocking buffer in +ENL tube was removed, and the beads were incubated with blocked phages transferred from -ENL tube for 30 minutes under slow rocking. Phages were then removed, and the beads were washed three times (or four times for the 3rd round of selection) with 1 mL of wash buffer. The affinity selection followed the same procedure as above for all proceeding steps. The enriched library from the ENL selection was analyzed by next generation sequencing.

**Characterization of Selected Peptide Inhibitors**

*AlphaScreen Assay to Validate ENL Inhibitors*:
His-ENL, His-AF9 (aa 1-149, EpiCypher), and biotinylated H3K27cr (aa 15-33, EpiCypher) were used to assay the synthesized inhibitors. The lyophilized peptide inhibitors were dissolved in DMF and serially diluted into assay buffer (50 mM HEPES, 100 mM NaCl, 0.1% BSA, 0.05% CHAPS, pH 7.4) and incubated in the presence of 120 nM His-tagged protein at 37 °C for 30 minutes. 12.5 µL of biotinylated H3K27cr (400 nM) was aliquoted into a light gray 384-well Alphaplate (Perkin Elmer) and incubated at 37 °C. 12.5 µL of the inhibitor/protein solution was added to the H3K27cr solution, and incubated for 30 minutes at 37 °C. 25 µL of AlphaScreen histidine (nickel chelate) detection beads (Perkin Elmer) was added at 20 µg/mL each bead (streptavidin donor beads and nickel chelate acceptor beads), and the plate was incubated at 37 °C for 30 minutes. The plate was cooled to room temperature for 10 minutes, and then the beads were excited at 680 nm and the luminescence at 615 nm was read.

*Biolayer Interferometry to Validate ENL Ligands*:
His-ENL, His-AF9 (aa 1-149, EpiCypher), His-GAS41 (aa 15-160, EpiCypher), and YEATS2-His (aa 200-345, EpiCypher) were reconstituted in water and diluted into assay buffer (20mM Tris, 300 mM NaCl, pH 7.9) for immobilization onto Octet Ni-NTA biosensors (Sartorius). Lyophilized peptides were dissolved in DMF and diluted to designated concentrations in assay buffer. Biolayer

interferometry assays were performed in 96 well plates (GreinerBio-One, polypropylene, flat-bottom) using an Octet Red96 System (Sartorius). Wells were filled with 200 μL of assay buffer, blocking buffer (assay buffer + 0.1% BSA), protein solution, peptide solution, regeneration solution, and reload solution. For immobilization, His-tagged proteins were immobilized onto Ni-NTA sensors at 20 μg/ml for 600 s. Sensors were then dipped into blocking buffer for 180 s, assay buffer for 300 s, peptide solution for 300 s, and back to assay buffer for 300 s. If multiple samples were tested, Ni-NTA sensors can be reconditioned in regeneration (10 mM glycine, pH 1.7) solution followed by reload solution (10 mM $NiCl_2$) for 30 and 60 s, respectively. Measurements were carried out at 30 °C. Data were analyzed within the ForteBio Data Analysis software. Data were processed by double subtracting ligand only and protein only reference wells and aligning the data to the beginning of the association. Then, global kinetic fit was performed for all sensogram curves with a 1:1 model. Steady state $K_d$ values were reported.

**Cell-Based Assays**

*Cell Line Culture*:

HEK 293T/17, MV4;11, and Jurkat cell lines were purchased from American Type Culture Collection. MOLM-13 cell line was purchased from AddexBio. MV4;11, Jurkat, and MOLM-13 cells were maintained in RPMI 1640 medium with 10% fetal bovine serum (FBS, Thermo Scientific), penicillin (100 IU/mL), and streptomycin (100 μg/mL). HEK 293T/17 cells were cultured in Gibco high glucose DMEM medium (Thermo Scientific) with 10% FBS. All cells were cultured at 37 ºC with 5% $CO_2$. Cells at logarithmic phase were used for following experiments.

*Cell Proliferation Assay*:

Cell proliferation assays were performed in 96-well tissue culture plates (Corning). The cells were seeded at 20,000 cells/well in 200 μL of growth medium with test compound (tENL-S1f at a final concentration of 10 μM) and control (DMSO at a final concentration of 0.1%) in triplicate. Cell numbers were determined every 3–4 days using the Countess automated cell counter (Invitrogen). After measurement, cell cultures were centrifuged (200 rcf, 5 min, room temperature), and pelleted cells were diluted in fresh medium and reseeded at 20,000 cells/well with the inhibitor. The cumulative cell count was acquired through back calculation.

*Preparation of ENL stably expressing cell line*:

The complementary DNA encoding ENL YEATS (aa 1-148) was cloned into pCDH-EF1-Nluc vector (Addgene, plasmid #73024) between sites *EcoRI* and *SalI* to afford pCDH-EF1-Nluc-ENL. The resulting plasmid was transformed into TOP10 *E. coli* cells for amplification and extracted by EndoFree Plasmid Midi Kits (Omega Bio-tek). For packaging lentivirus particles, HEK293T/17 cells were grown to 70-80% confluency and then co-transfected with three plasmid, pCDH-EF1-Nluc-ENL, psPAX2 (Addgene, plasmid #12260), and PMD2.G (Addgene, plasmid #12259) using polyethyleneimine. The transfected cells were maintained in DMEM for two days, and the viral particles were harvested from the supernatant. The lentivirus supernatant was centrifuged (500 rcf, 5 min, 4 ºC), filtered through a 0.45 μm PES filter to remove cell debris. The solution was aliquoted and stored at -80 ºC. For transduction, the lentivirus solution was added to HEK293/17 cells (MOI 3-5) and incubated for 24 h with 4 μg/mL polybrene (Sigma-Aldrich) before the medium was refreshed. After two days, the cells were transferred to medium with puromycin to select transduced cells.

*Fluorescence Polarization*:
Tracer 1 (10 nM) and varying concentrations of His-ENL (14 nM to 7.2 µM) were mixed in a 200 µL of assay buffer (50 mM HEPES, 100 mM NaCl, 0.01% Tween20, pH: 7.4). The solutions were incubated in black 96-well plates and then measured fluorescence polarization in a microplate reader at Ex/Em = 530 nm/590 nm.

*NanoBRET Assay to Validate ENL Inhibitors*:
HEK293T/17 cells harboring pCDH-EF1-Nluc-ENL were cultivated in growth medium overnight at 37 °C and 5% $CO_2$. Cells were harvested and resuspended in assay medium (phenol red-free OptiMEM I reduced serum medium, Thermo Scientific) at 2 x $10^5$ cells/mL. Cell suspension (85 µL/well) was added to 96-well culture plate (Corning). To cells were added 5 µL of 20X Tracer 1 solution (50 µM for validating ENL-S1 peptides; 5-50 µM for profiling tENL-S1f) prediluted with Tracer Dilution Buffer (Promega) at a 1:4 ratio, per the manufacturer's instructions, and 5 µL of 20% DMSO in Tracer Dilution Buffer as no tracer control. Following a quick spin (100 rcf, 1 min) to ensure reagent mixing, the cells were treated with 10 µL of 10X test compounds (prepared and serially diluted in DMSO; prediluted 100-fold with assay medium) and control (1% DMSO in assay medium). The mixtures were incubated at 37 °C and 5% $CO_2$ for 2 h before being equilibrated to room temperature for 15 min. The samples were added with 50 µL of 3X NanoBRET Nano-Glo Substrate plus Extracellular NanoLuc Inhibitor Solution (1:166 dilution of NanoBRET Nano-Glo Substrate plus 1:500 dilution of Extracellular NanoLuc Inhibitor in phenol red-free Opti-MEM assay medium). Following a quick spin (100 rcf, 1 min), a plate reader equipped with the LUM 610 LP 450 module was used to measure the emissions at 450 nm (donor) and 610 nm (acceptor).

The final BRET ratio was calculated by dividing the acceptor emission (610 nm) by the donor emission (450 nm). The BRET ratio was calculated using the following equation:
BRET ration = $[(\text{Acceptor}_{\text{sample}} \div \text{Donor}_{\text{sample}}) - (\text{Acceptor}_{\text{no tracer}} \div \text{Donor}_{\text{no tracer}})] \times 1{,}000$

*qRT-PCR*:
MOLM-13 cells were incubated with test compound (tENL-S1f at a final concentration of 10 and 50 µM) and control (DMSO at a final concentration of 0.1%) for 24 hours. The total RNA was extracted using RNeasy Mini plus Kit (Qiagen) and reverse-transcribed using the SuperScript VILO cDNA Synthesis Kit (Invitrogen) at 100 ng/µL RNA. Quantitative real-time PCR (qRT-PCR) analyses were performed in triplicate using SYBR Select Master Mix (Applied Biosystems) and the Bio-Rad CFX96 real-time PCR detection system with the primer pairs listed in Table S1. Gene expressions were calculated following normalization to B2M levels using the comparative $C_t$ (cycle threshold) method.

Table S1. Primers used in qRT-PCR

| Gene | Sequence 5'-3' |
|---|---|
| HOXA9 | F: TACGTGGACTCGTTCCTGCT |
| | R: CGTCGCCTTGGACTGGAAG |
| MEIS1 | F: CCAGCATCTAACACACCCTTAC |
| | R: TATGTTGCTGACCGTCCATTAC |
| MYB | F: CTCCAAGAACTCCTACACCATTC |
| | R: GTCATCTGCTCCTCCATCTTTC |
| MYC | F: CACCGAGTCGTAGTCGAGGT |
| | R: TTTCGGGTAGTGGAAAACCA |

**Molecular Dynamic Simulations**

*Background*:
The interactions between tENL-S1f and YEATS proteins were predicted by simulations performed based on crystal structure PDB 5J9S (ENL YEATS) and 4TMP (AF9 YEATS). The Maestro (Schrödinger Release 2020-1: Maestro, Schrödinger, LLC, New York, NY, 2020) and Desmond (Schrödinger Release 2020-1: Desmond Molecular Dynamics System, D. E. Shaw Research, New York, NY, 2020; Maestro-Desmond Interoperability Tools, Schrödinger, New York, NY, 2020) software was used for all simulations. Each of the simulations for investigating the protein–peptide interactions followed the steps described below.

*Protein Preparation*:
The structures of ENL and AF9 YEATS with native substrate were available in the PDB. To simulate the interactions between tENL-S1f and YEATS proteins, PDB 5J9S and 4TMP crystal structures were chosen as templates. Both structures showed high resolution and residue completeness, and contained a native acetyl-lysine peptide that were modified into tENL-S1f. The Protein Preparation Wizard (Schrödinger Release 2020-1: Protein Preparation Wizard; Epik, Schrödinger, LLC, New York, NY, 2020; Impact, Schrödinger, LLC, New York, NY; Prime, Schrödinger, LLC, New York, NY, 2020) was used to prepare both 5J9S and 4TMP structures for simulation at a pH of 7.5 ± 0.[4] Crystal structure waters were maintained. The H-bonding network was optimized using the PROPKA tool at a pH of 7.5 after initial preparation, water molecule orientations were sampled, and a restrained minimization was performed on all atoms using the OPLS3 force field.[5]

*Building Molecular Dynamics Simulations Model*:
The System Builder tool included in the Maestro interface distribution inside Desmond Module was used to prepare the model for a Desmond MD simulation. The native acetyl-lysine peptide was modified to tENL-S1f for both crystals before applying the solvent box. An orthorhombic solvent box was constructed with a distance of 10.0 Å around the inhibitor–protein binding channel. The solvent box was size-minimized before applying explicit water molecules to the system via TIP4P water model[6-7] while OPLS3[5] force field for all other atoms. The protein–inhibitor assembly was charge neutralized using a physiological NaCl concentration of 0.15 M.

*Molecular dynamics simulations with Desmond*:

Both inhibitor-protein assembly were simulated with The Desmond multisim molecular dynamics protocol[8] using NPT ensemble (1.0135 bar, 310.15 K) for a 10-ns production run taking a snapshot every 10 ps, resulting in a trajectory with 1000 frames. A relaxation protocol was used for both crystals before the production run, as implemented in the Desmond Schrödinger software Maestro interface, and outlined as follows: (1) restrained minimization steps. (2) Brownian dynamics NVT, T = 10 K, small timesteps, and restraints on solute-heavy atoms, 100 ps. (3) NVT, T = 10 K, small timesteps, and restraints on solute-heavy atoms, 12 ps. (4) NPT, T = 10 K, and restraints on solute-heavy atoms, 12 ps. (5) NPT and restraints on solute-heavy atoms, 12 ps. (6) NPT and no restraints, 24 ps. (7) Production run.

*Protein–inhibitor interactions*:

Both simulation results were imported, and ligand interaction tables were then generated using the ligand interaction diagram module in the Desmond distributed Schrödinger software. Key protein–ligand interactions formed between YEATS proteins and tENL-S1f, such as pi-pi stacking and hydrogen-bond interactions were reported (Figure 6 and S5).

#### Supplementary Figures

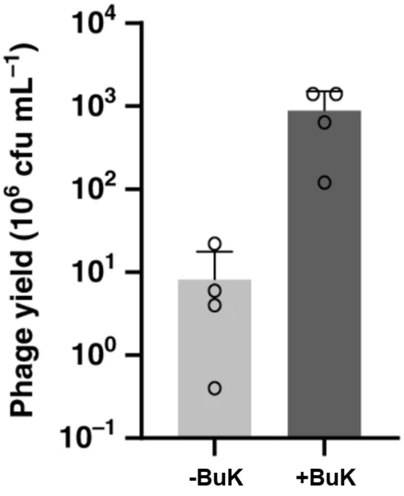


Figure S1. Phage yield in the presence and absence of BuK. The yield is displayed in millions of colony-forming units. Error bars represent one standard deviation of the mean of four independent experiments (n = 4).

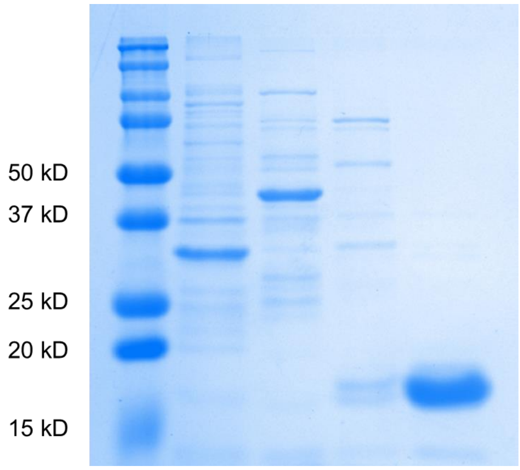


Figure S2. Expression of Avi-SUMO. Expected molecular weight: 15.3 kDa. Lane 1: Flow-through after incubation with Ni-NTA resin, Lane 2 and 3: Fractions collected from washing steps, Lane 4: Purified Avi-SUMO.

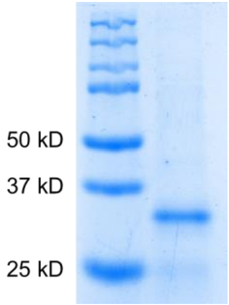


Figure S3. Expression of Avi-SUMO-ENL. Expected molecular weight: 32.3 kDa.

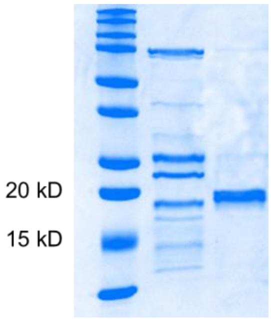


Figure S4. Expression of His-ENL. Expected molecular weight: 20.0 kDa. Lane 1: Fraction collected from washing step, Lane 2: Purified His-ENL.

| | | A | B | C | D | E | F | G | H | I | J |
|---|---|---|---|---|---|---|---|---|---|---|---|
| | 1 | | V1 | V2 | V3 | V4 | V5 | V6 | V7 | n | enrichment |
| | 2 | | | | | | | | | | |
| ENL-S1 | 3 | 1 | Y | D | V | Y | C | Y | TAG | 62878 | 451.25532 |
| ENL-S2 | 4 | 2 | W | W | I | I | E | TAG | G | 35160 | 180.87386 |
| | 5 | 20 | Q | C | G | P | R | TAG | D | 9329 | 8.6249455 |
| | 6 | 13 | H | L | T | L | F | TAG | G | 2944 | 15.843725 |
| | 7 | 5 | Y | L | Y | TAG | V | P | C | 1065 | 133.05179 |
| | 8 | 25 | H | A | I | Y | C | Y | TAG | 786 | 6.5139746 |
| | 9 | 17 | H | Y | V | L | F | TAG | G | 640 | 11.719518 |
| | 10 | 4 | Y | D | V | Y | C | Y | Y | 357 | 133.80701 |
| | 11 | 3 | W | R | I | I | E | TAG | G | 187 | 140.2264 |
| | 12 | 9 | W | W | I | I | E | Q | G | 161 | 59.79532 |
| | 13 | 32 | R | T | V | Y | C | Y | TAG | 146 | 4.0119236 |
| | 14 | 16 | TAG | C | G | P | R | TAG | D | 125 | 12.486096 |
| | 15 | 6 | W | W | I | I | V | TAG | G | 99 | 73.766915 |
| | 16 | 7 | W | W | I | I | D | TAG | G | 94 | 69.990808 |
| | 17 | 8 | W | G | I | I | E | TAG | G | 91 | 67.725144 |
| | 18 | 50 | S | W | Y | I | T | F | TAG | 81 | -0.3422266 |
| | 19 | 53 | C | W | V | C | TAG | P | H | 80 | -0.5684449 |
| | 20 | 41 | W | W | V | A | G | P | TAG | 71 | -0.0070238 |

Figure S5. Next-generation sequencing data for the selection against ENL YEATS. Sequences were sorted by abundance. The enrichment was calculated by the (%R3-%R1)/%R1.

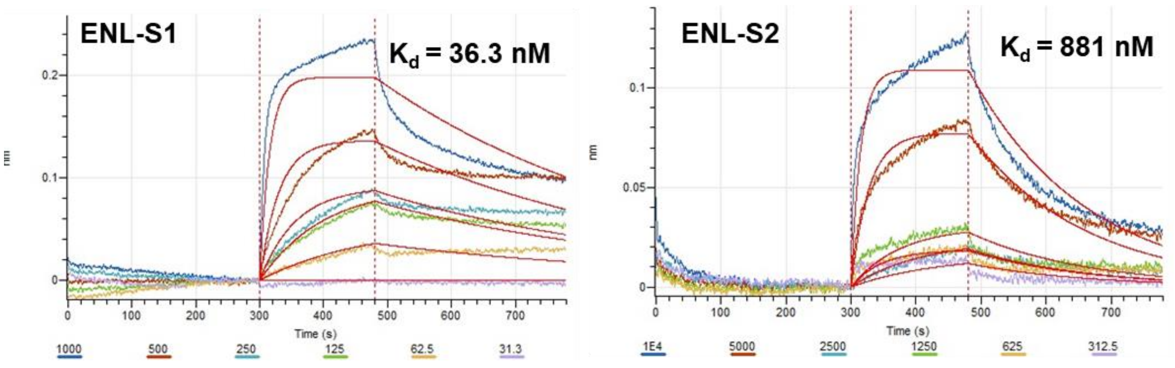


Figure S6. Biolayer interferometry (BLI) analysis of the selected peptides against ENL YEATS. Concentrations tested are listed below the curves for the corresponding peptides, along with $K_d$ values.

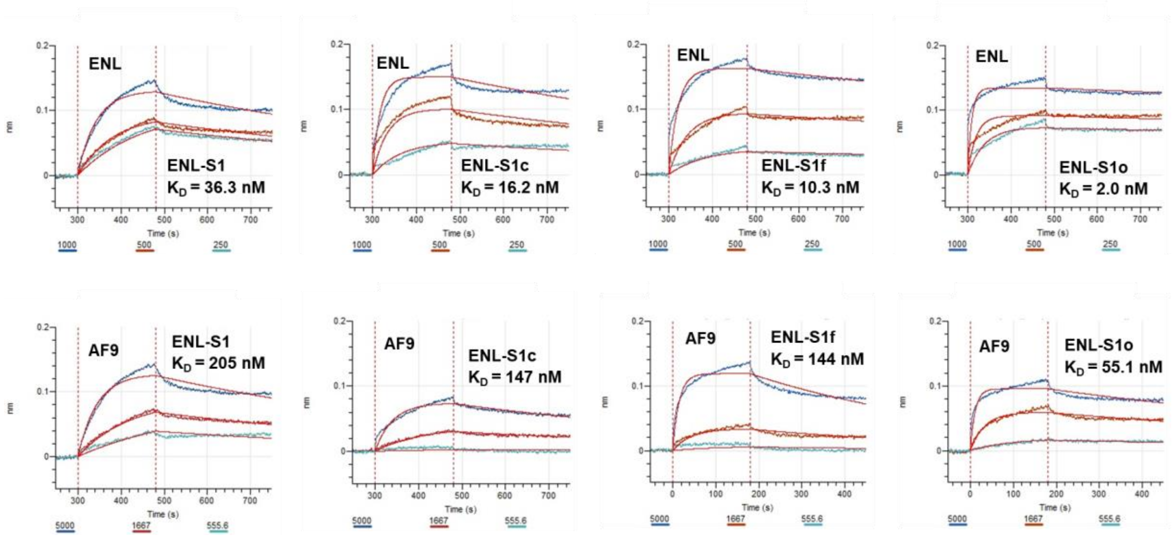


Figure S7. BLI analysis of **ENL-S1** derivatized peptides against ENL (Top row) and AF9 YEATS (Bottom row). Concentrations tested are listed below the curves for the corresponding peptides, along with $K_d$ values.

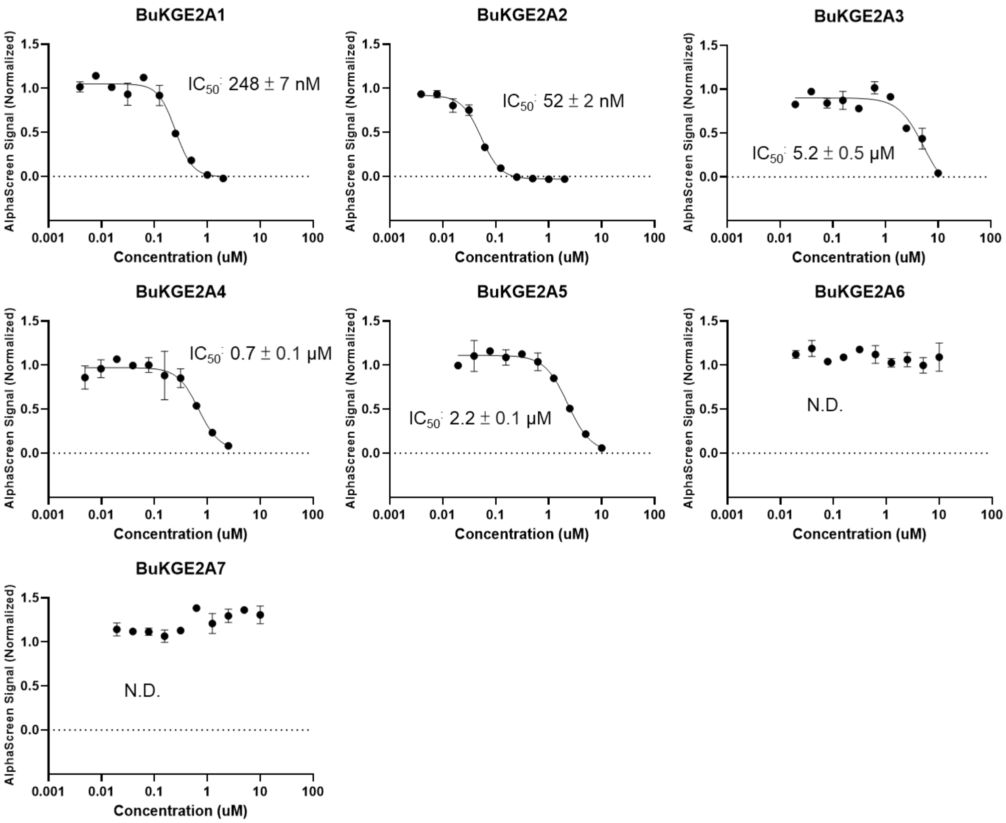


Figure S8. Alanine scan of **ENL-S1** measured with AlphaScreen. Data points and $IC_{50}$ values are given as the mean ± SD, n = 3

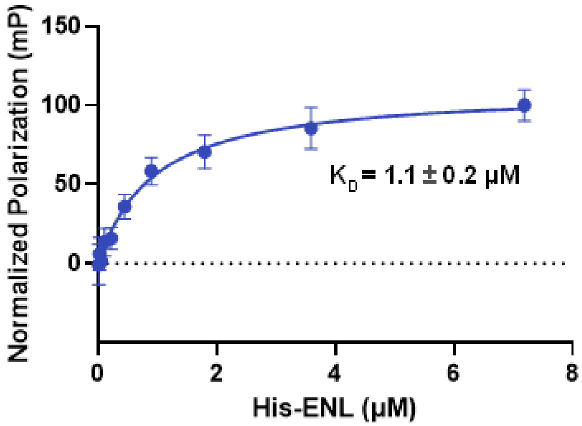


Figure S9. Fluorescence polarization analysis of Tracer1. Data points and $K_d$ value are given as the mean ± SD, n = 3

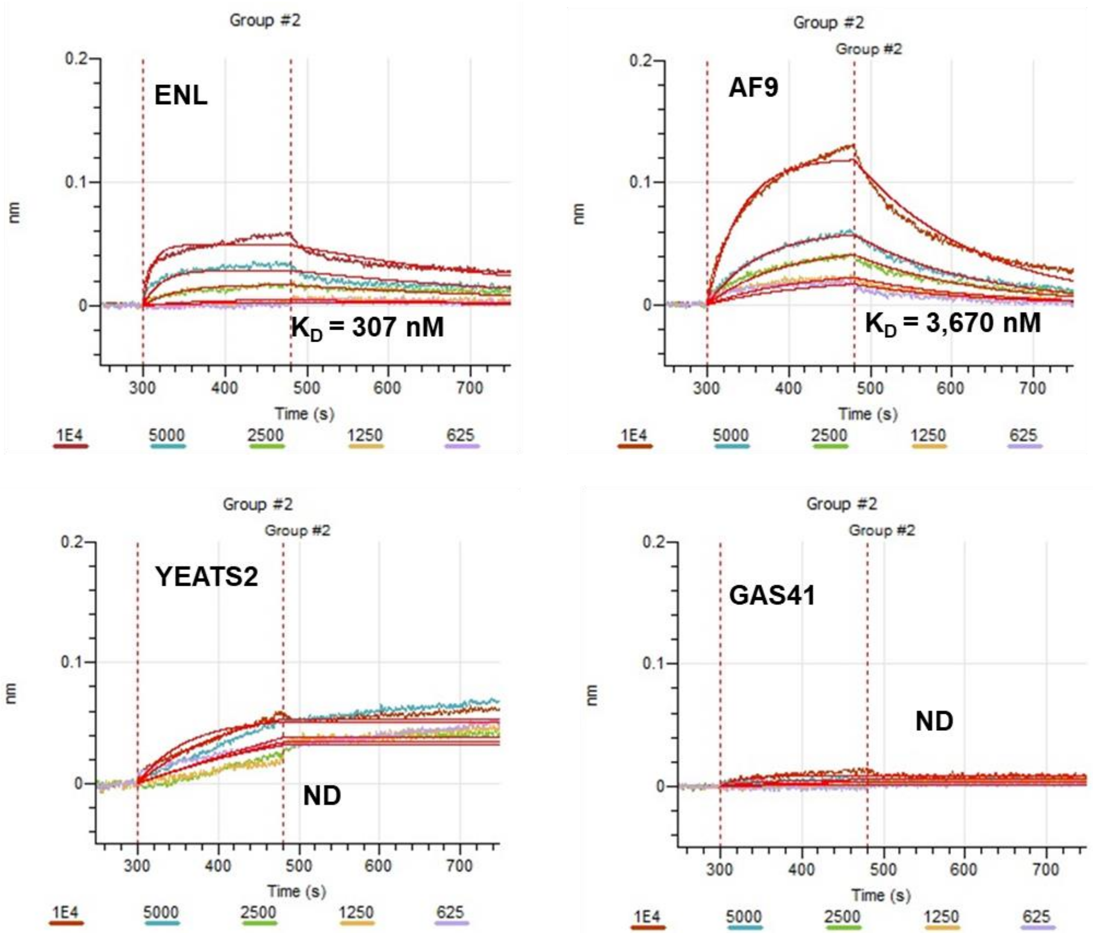


Figure S10. BLI analysis of **tENL-S1f** against four human YEATS proteins. Concentrations tested are listed below the curves for the corresponding peptides, along with $K_d$ values.

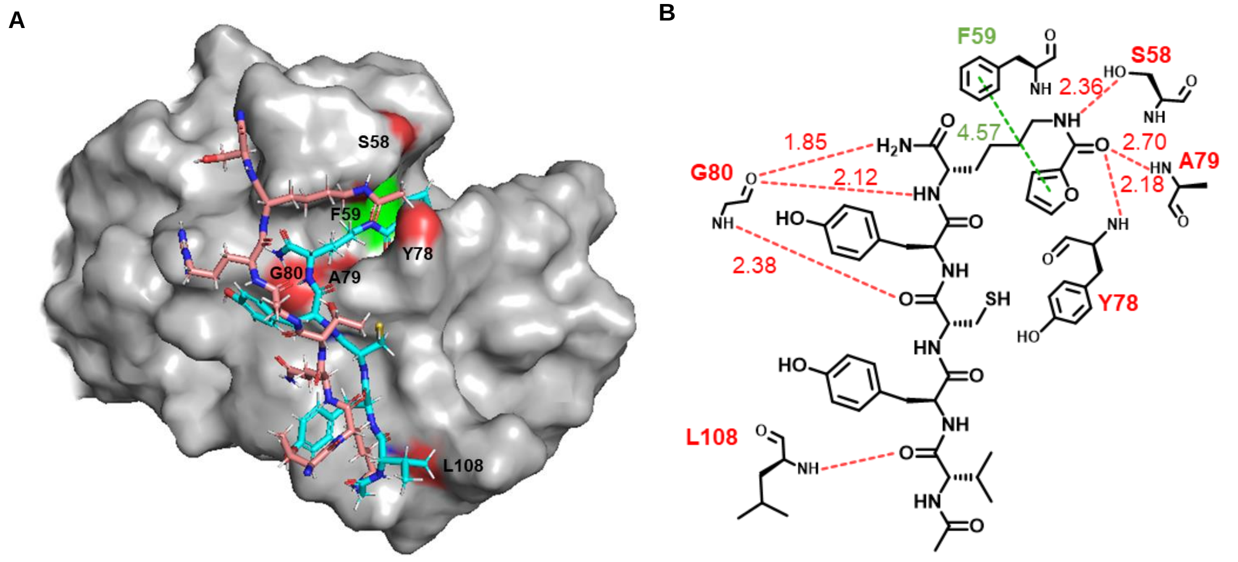


Figure S11. Molecular dynamic simulations to predict the interactions between **tENL-S1f** and AF9 YEATS. (A) The MD simulation predicted complex structure of **tENL-S1f** (cyan) bound to the substrate binding site of AF9 YEATS (gray). The native substrate H3K9Ac peptide is shown in salmon. (B) Interactions between **tENL-S1f** and AF9 YEATS predicted by molecular dynamic simulations. π-stacking (green) and hydrogen-bond (red) interactions are shown in dash lines with distance indicated in angstrom. MD simulations were performed based on the reported crystal structure (4TMP).

#### LC/MS of Synthesized Peptides

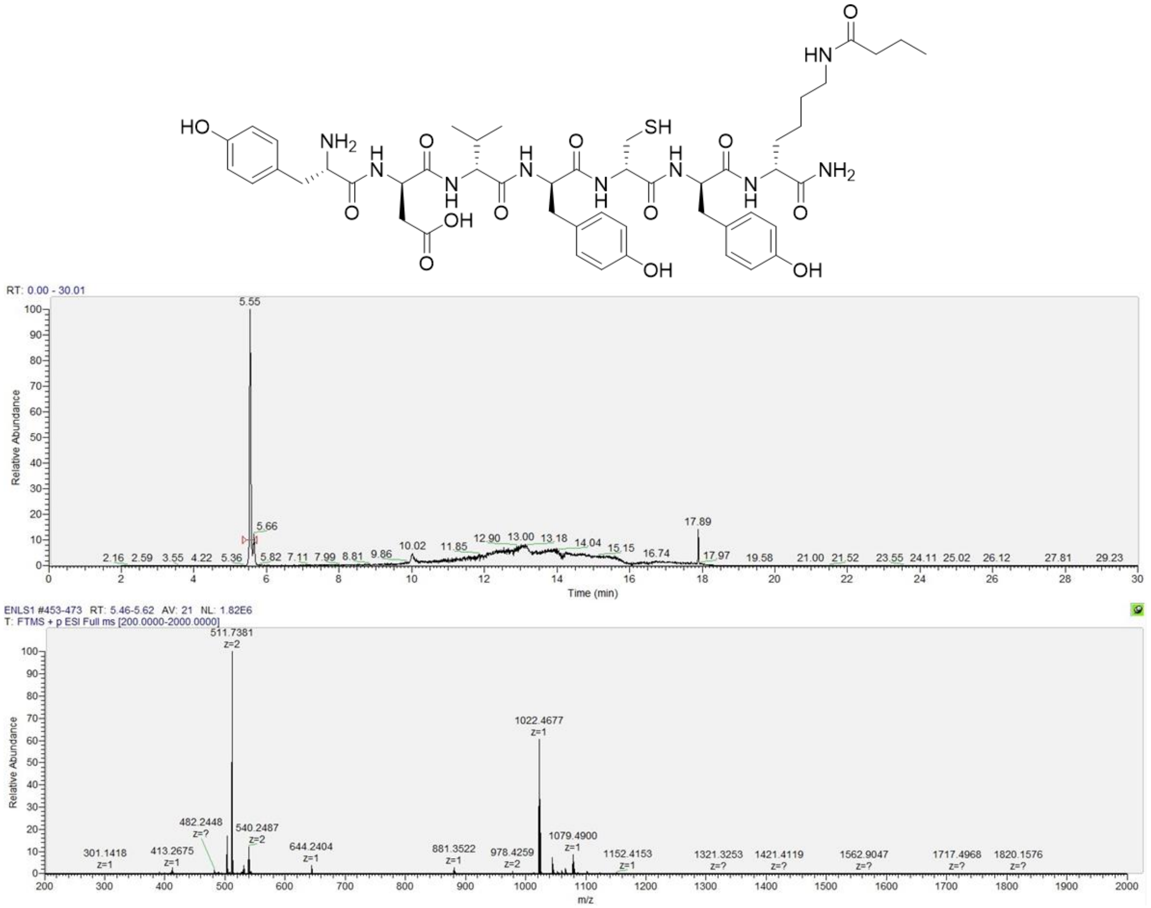


Figure S12. LC/MS of ENL-S1. Calculated $[M+H]^+$: 1022.4657 Da.

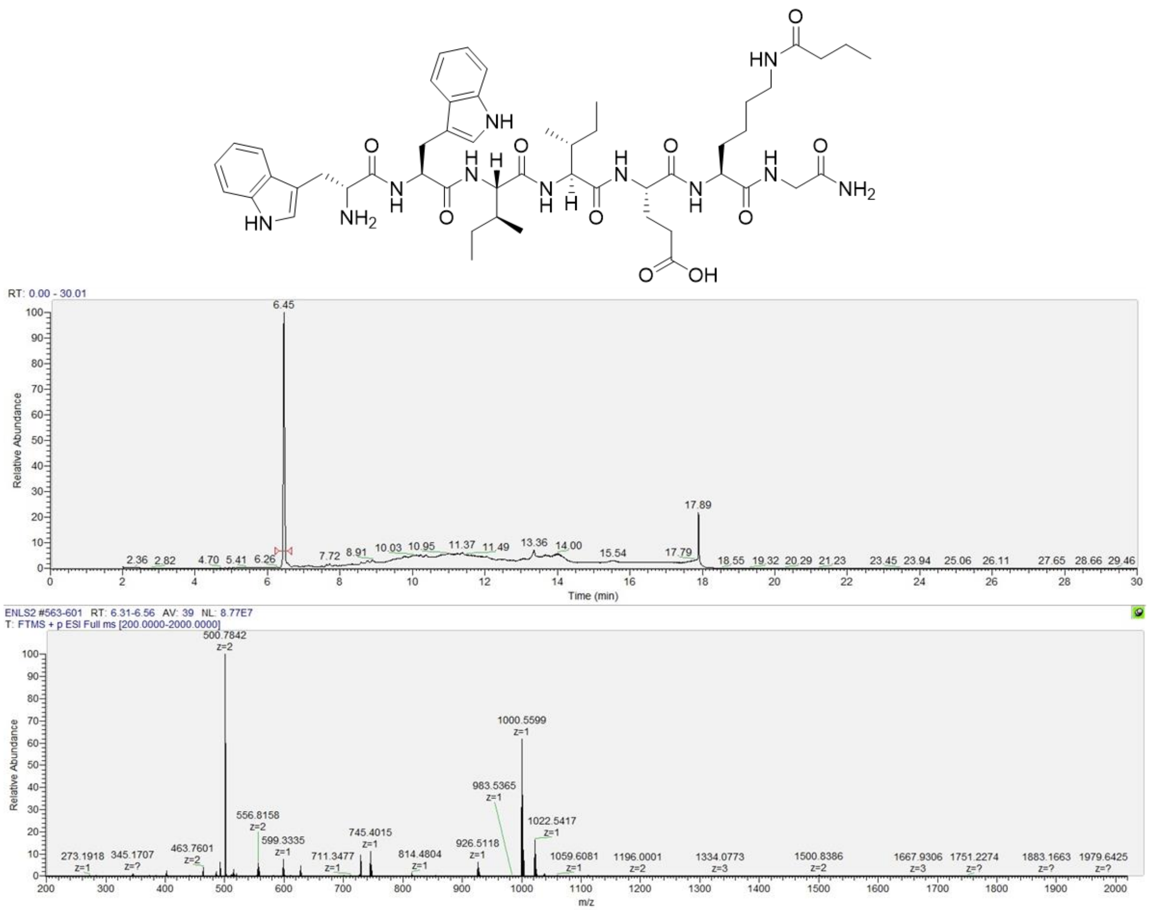


Figure S13. LC/MS of ENL-S2. Calculated $[M+H]^+$: 1000.5620 Da.

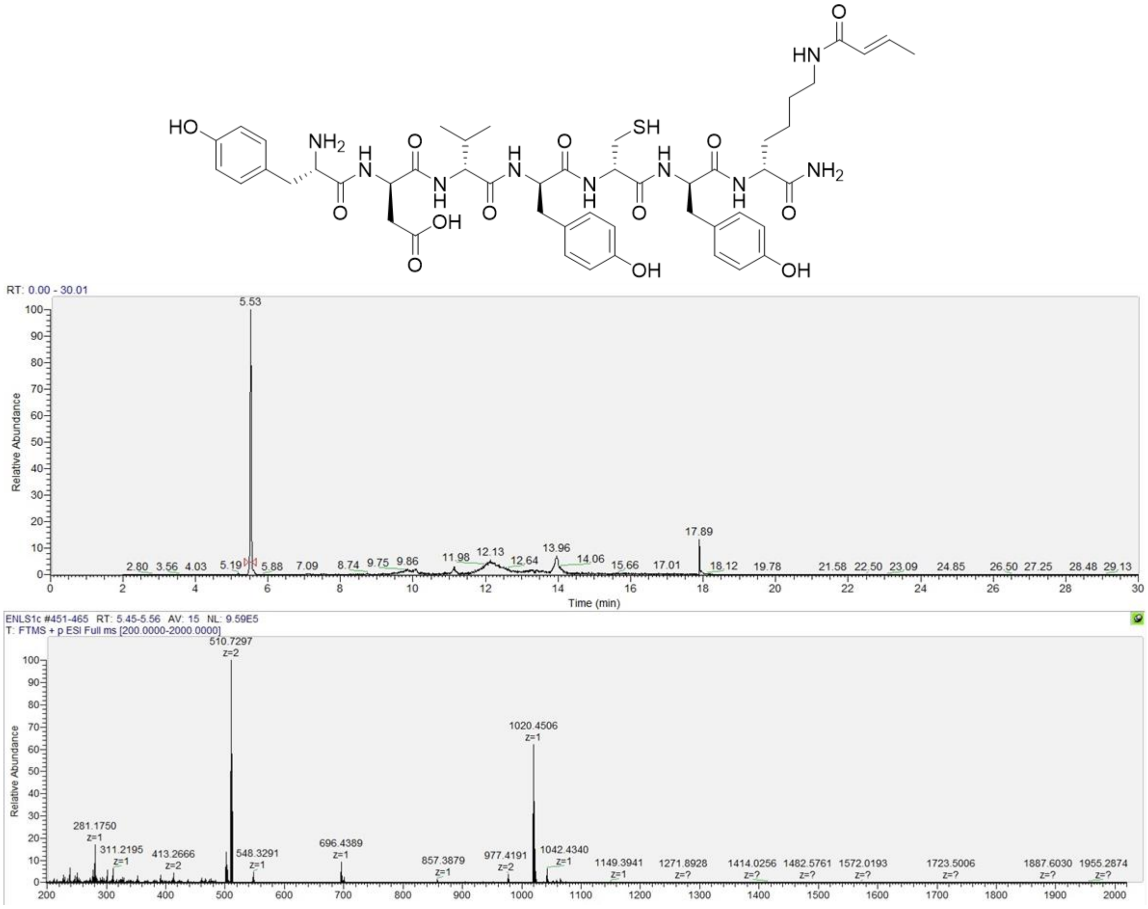

Figure S14. LC/MS of ENL-S1c. Calculated [M+H]$^{+}$: 1020.4501 Da.

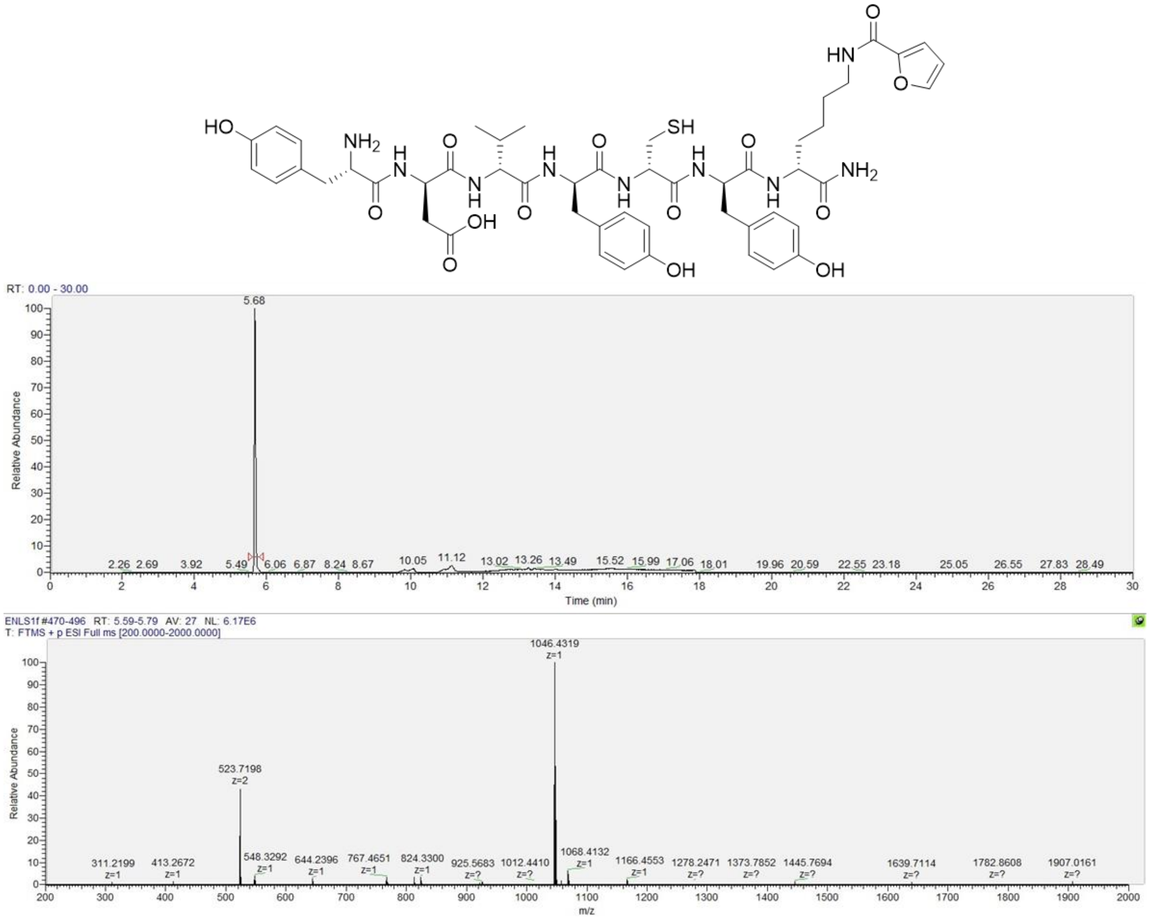


Figure S15. LC/MS of ENL-S1f. Calculated $[M+H]^+$: 1046.4293 Da.

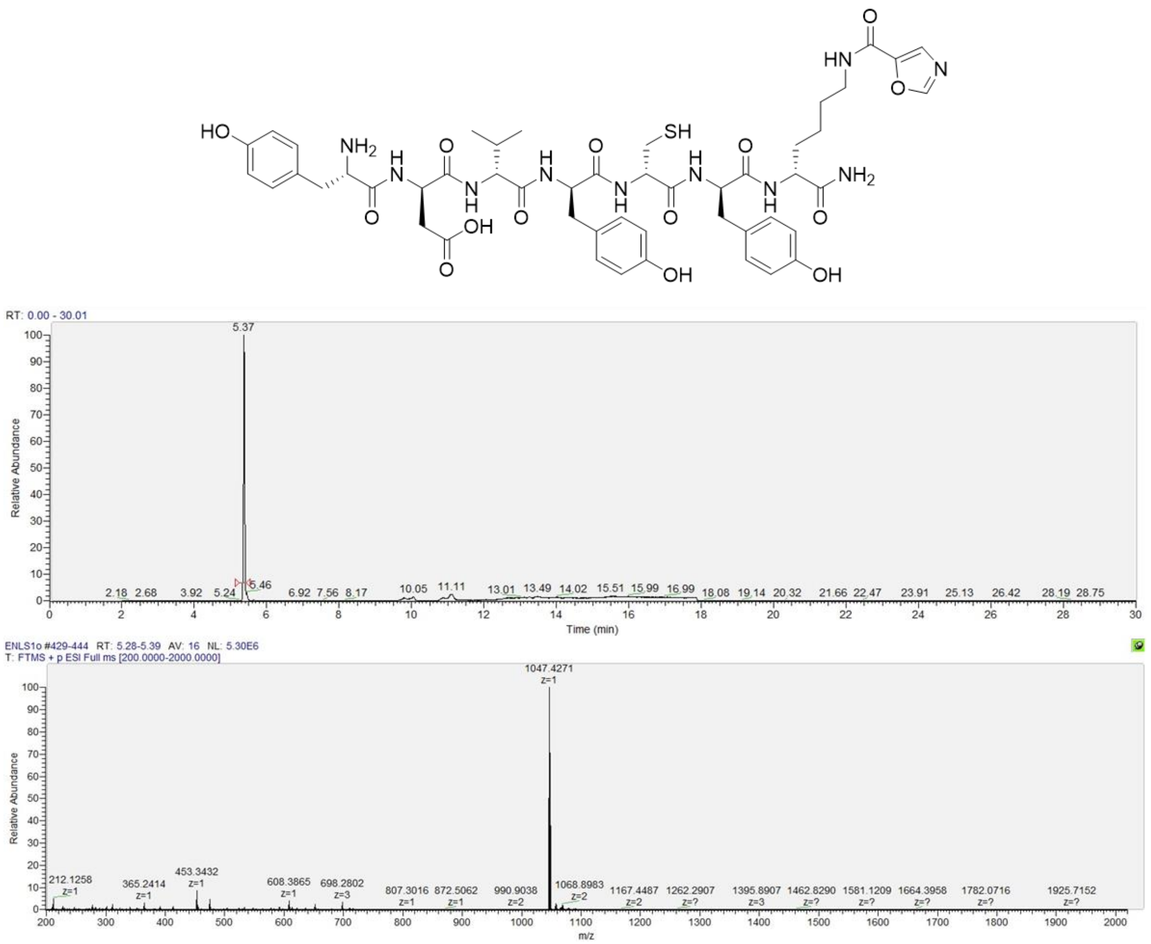

Figure S16. LC/MS of ENL-S1o. Calculated $[M+H]^+$: 1047.4246 Da.

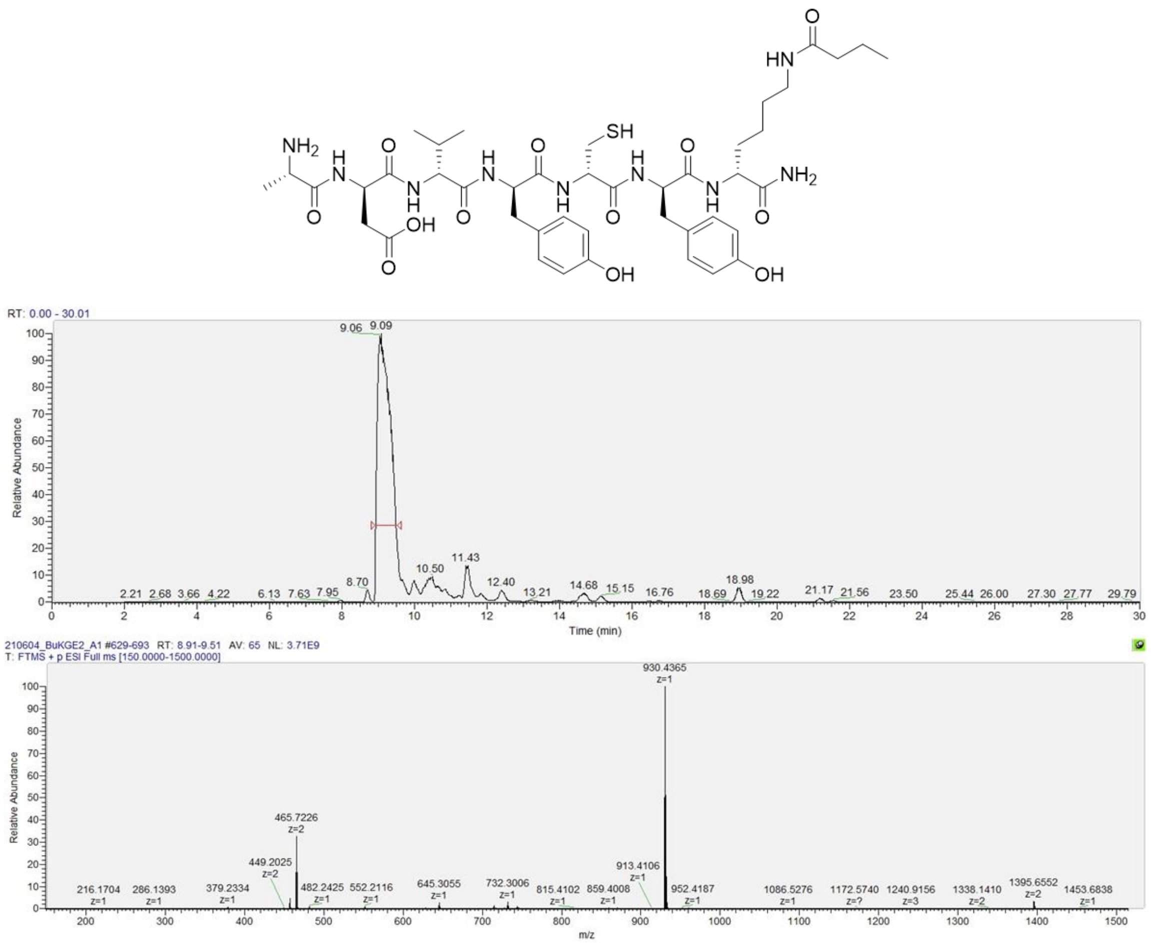


Figure S17. LC/MS of ENL-S1A1. Calculated $[M+H]^+$: 930.4395 Da.

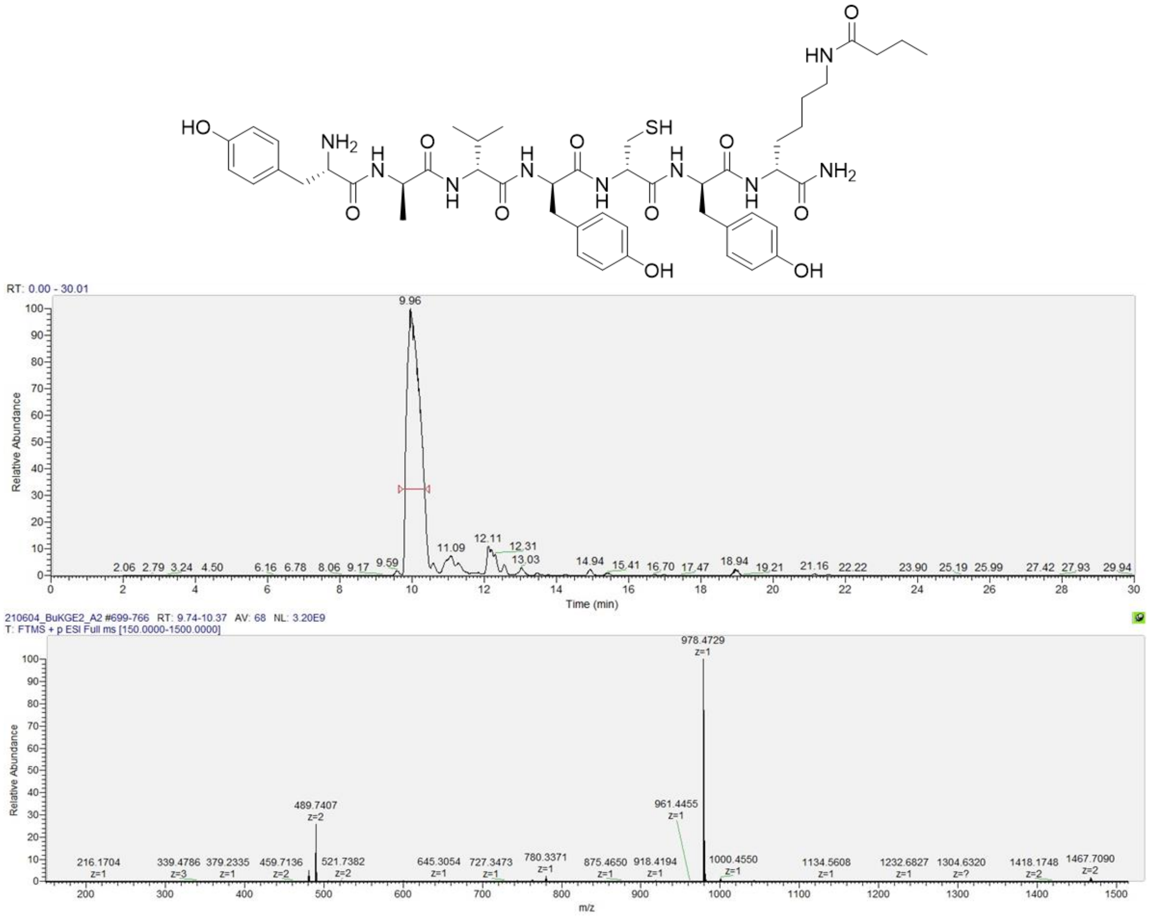


Figure S18. LC/MS of ENL-S1A2. Calculated $[M+H]^+$: 978.4759 Da.

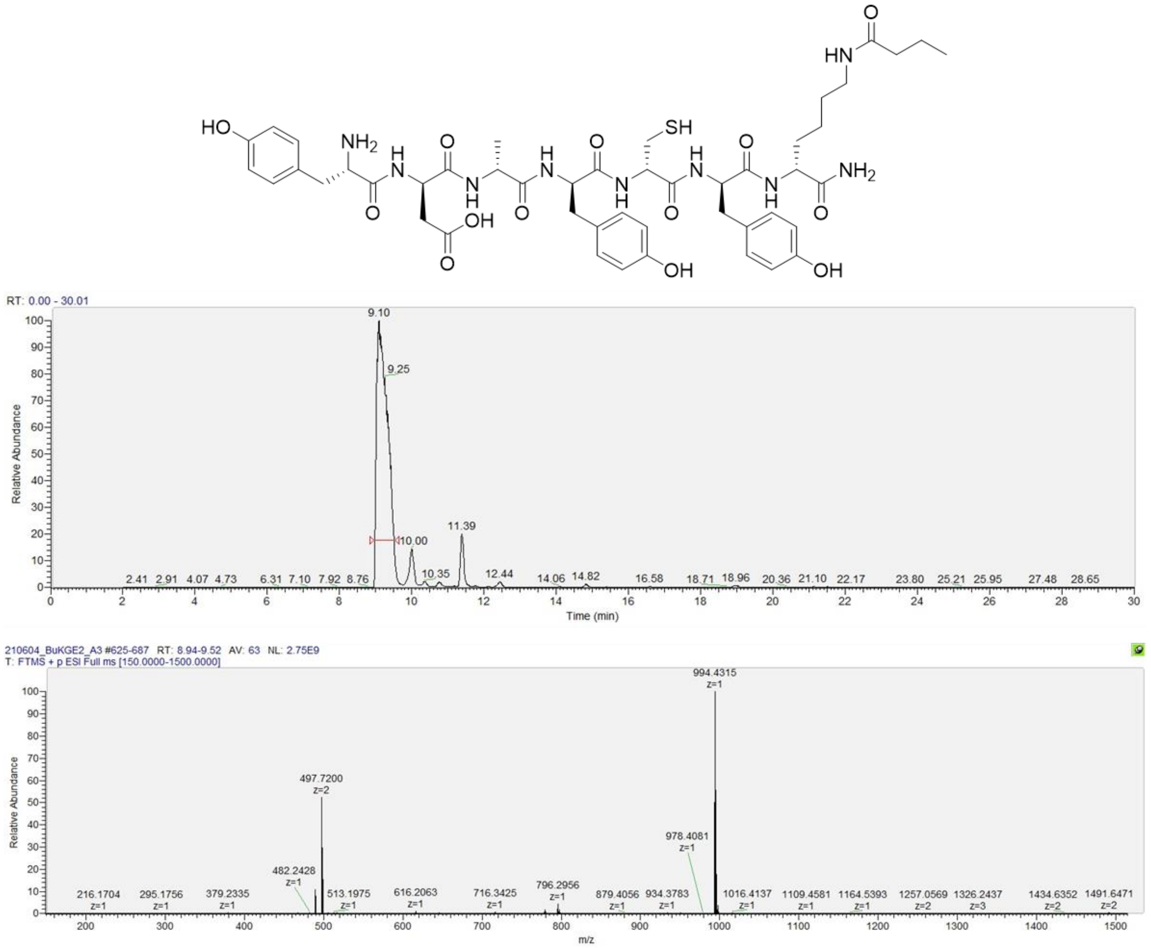

Figure S19. LC/MS of ENL-S1A3. Calculated $[M+H]^+$: 994.4344 Da.

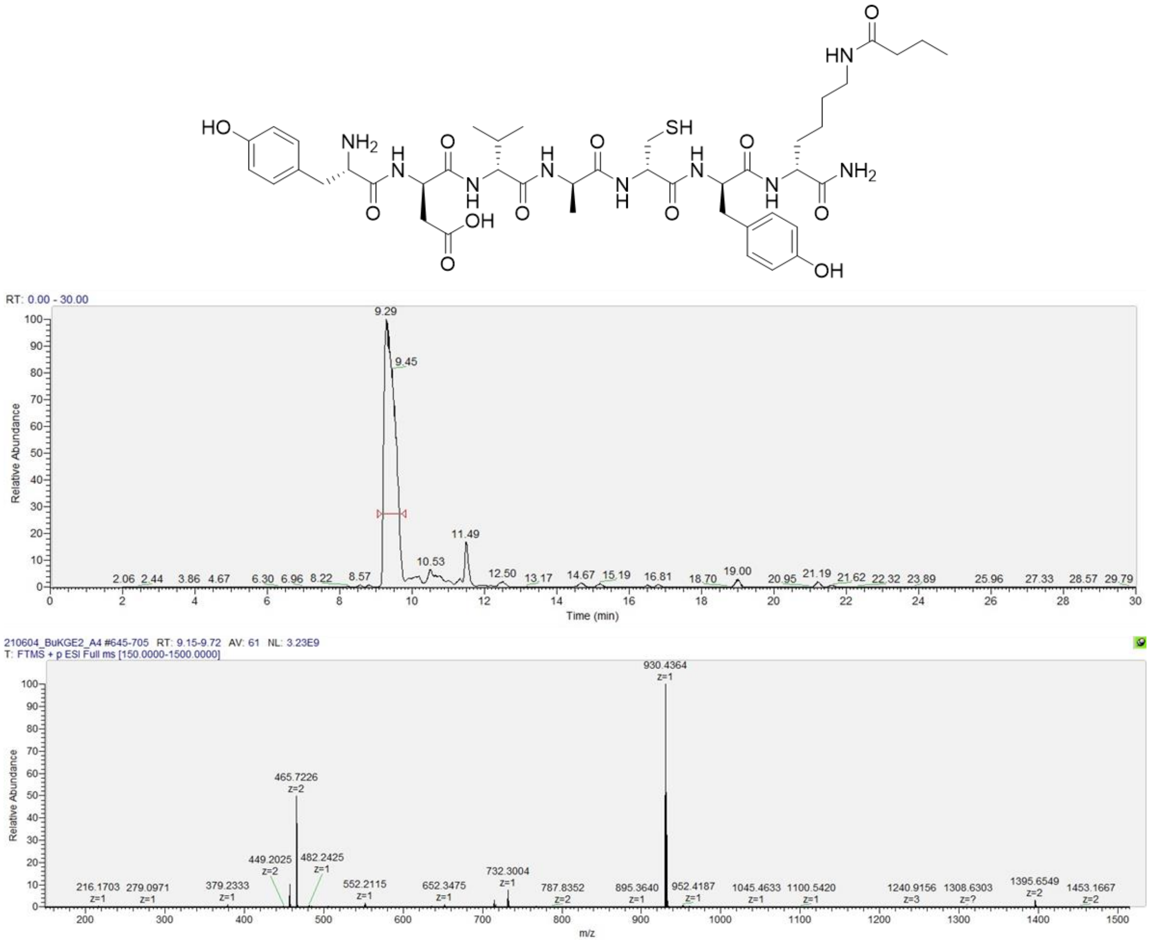


Figure S20. LC/MS of ENL-S1A4. Calculated $[M+H]^+$: 930.4395 Da.

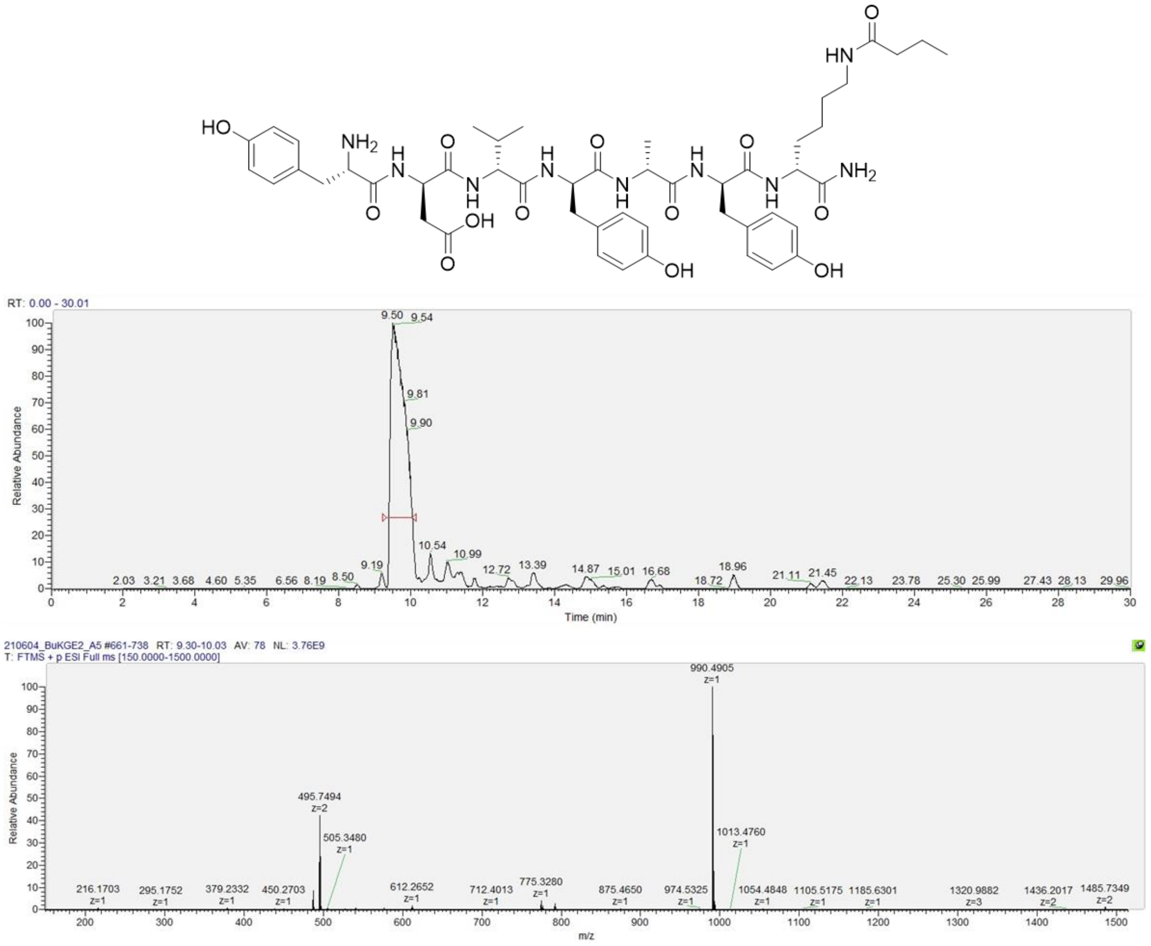


Figure S21. LC/MS of ENL-S1A5. Calculated $[M+H]^+$: 990.4937 Da.

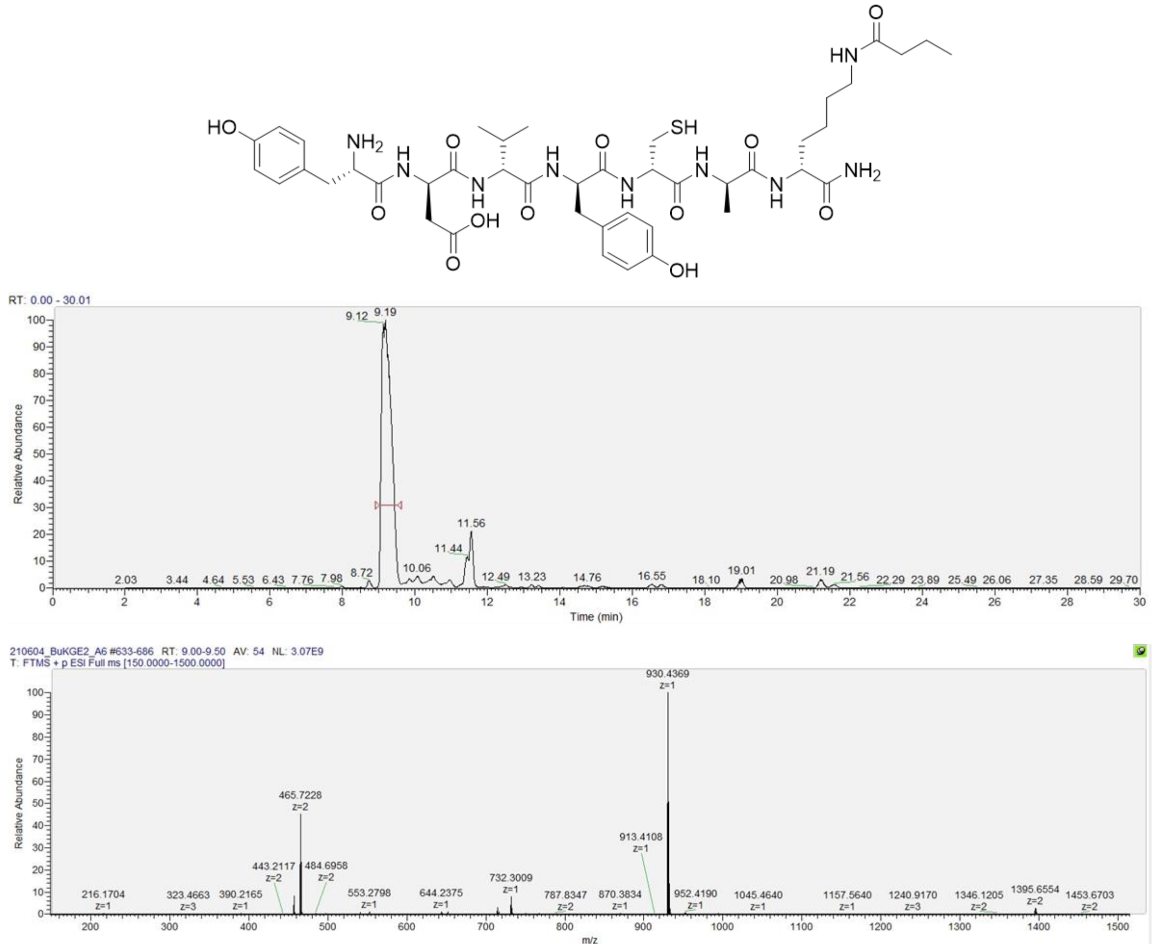


Figure S22. LC/MS of ENL-S1A6. Calculated $[M+H]^+$: 930.4395 Da.

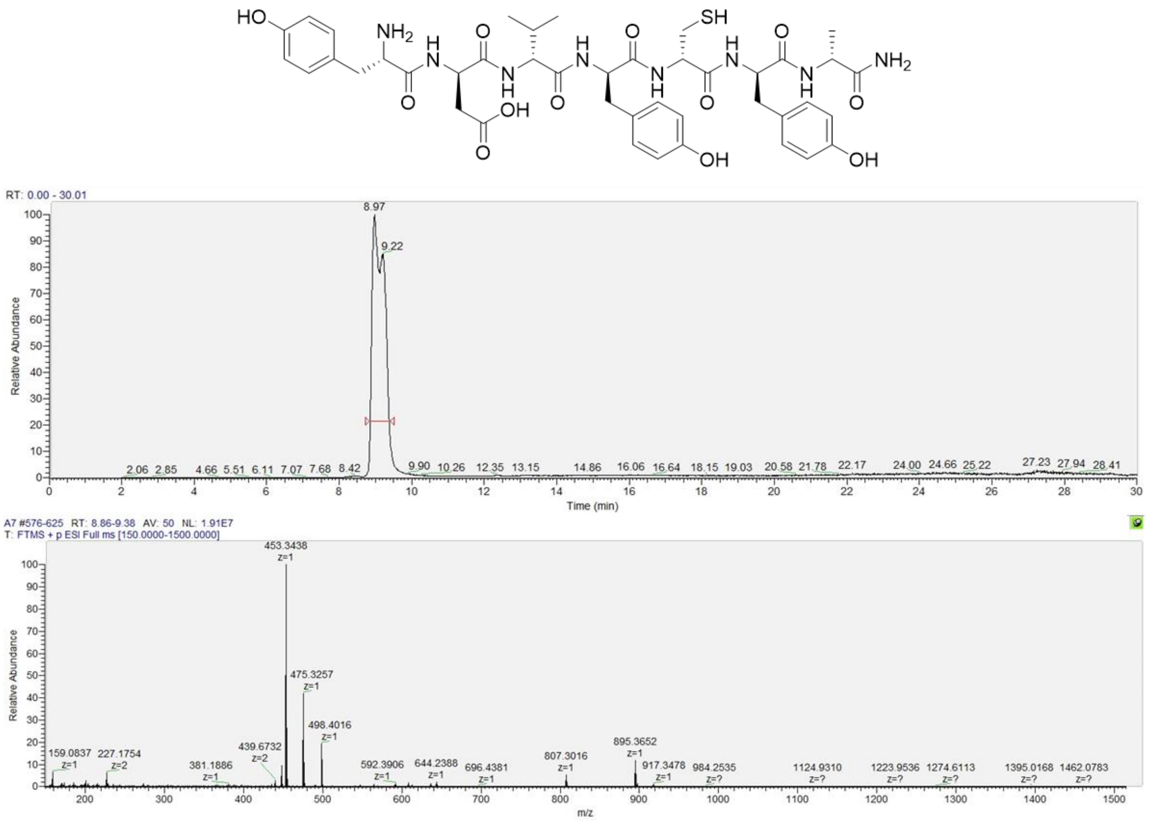


Figure S23. LC/MS of ENL-S1A7. Calculated $[M+H]^+$: 895.3660 Da.

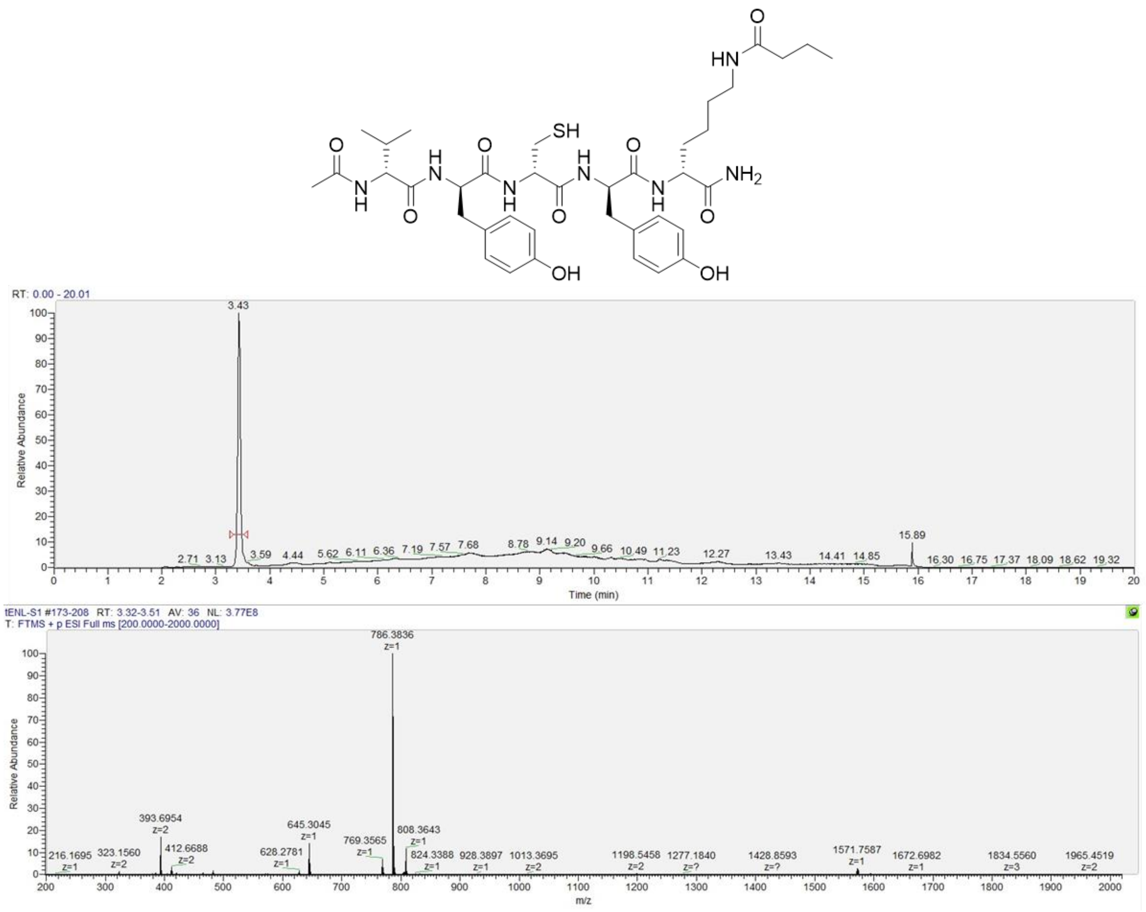


Figure S24. LC/MS of tENL-S1. Calculated $[M+H]^+$: 786.3860 Da.

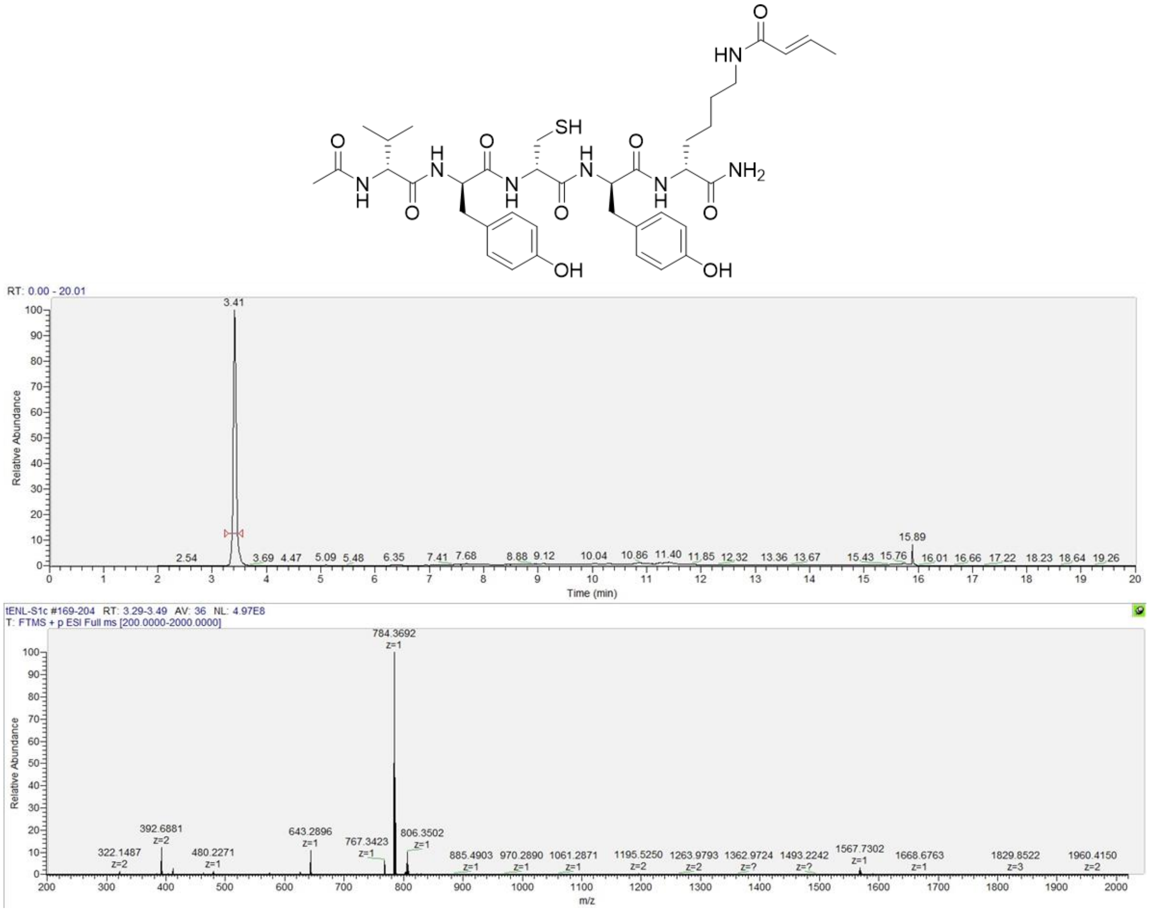


Figure S25. LC/MS of tENL-S1c. Calculated [M+H]$^+$: 784.3704 Da.

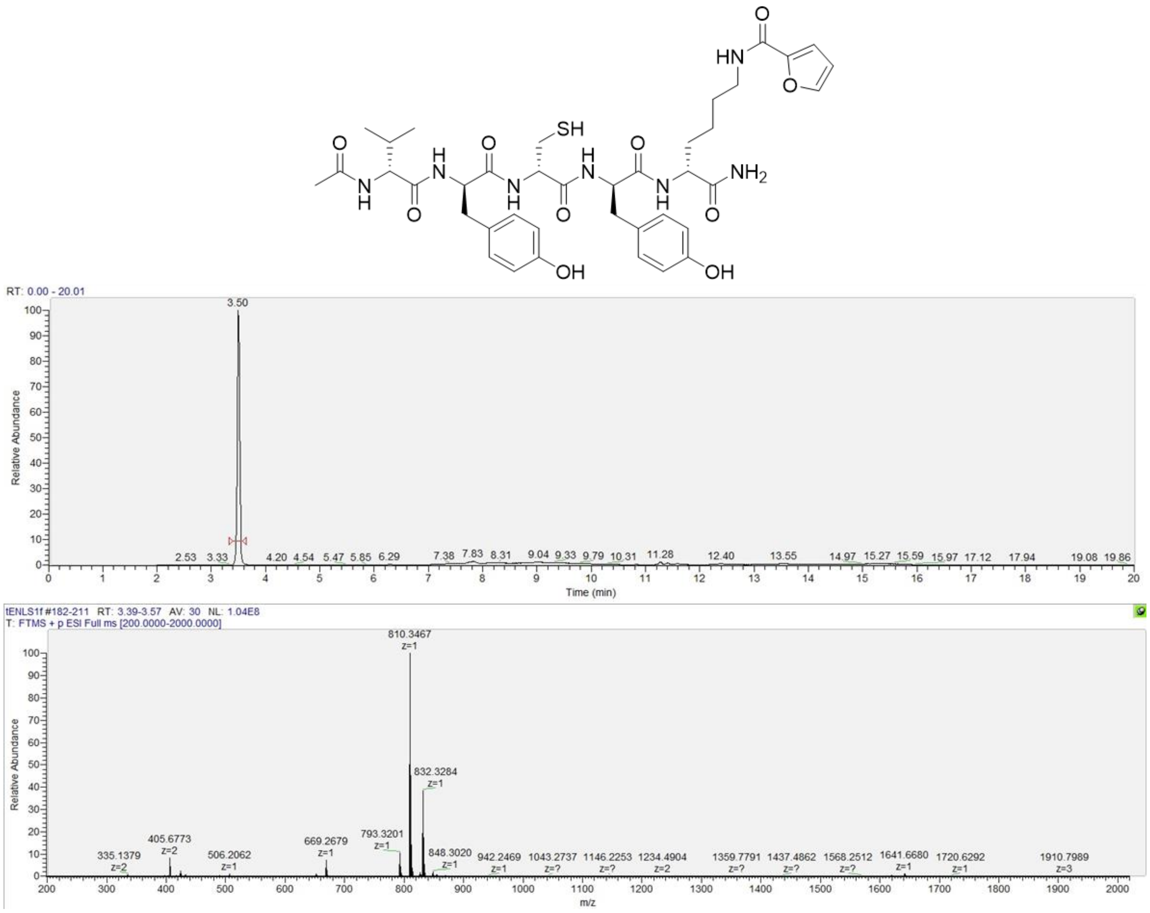

Figure S26. LC/MS of tENL-S1f. Calculated $[M+H]^+$: 810.3496 Da.

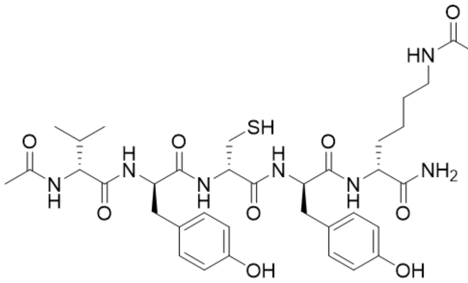

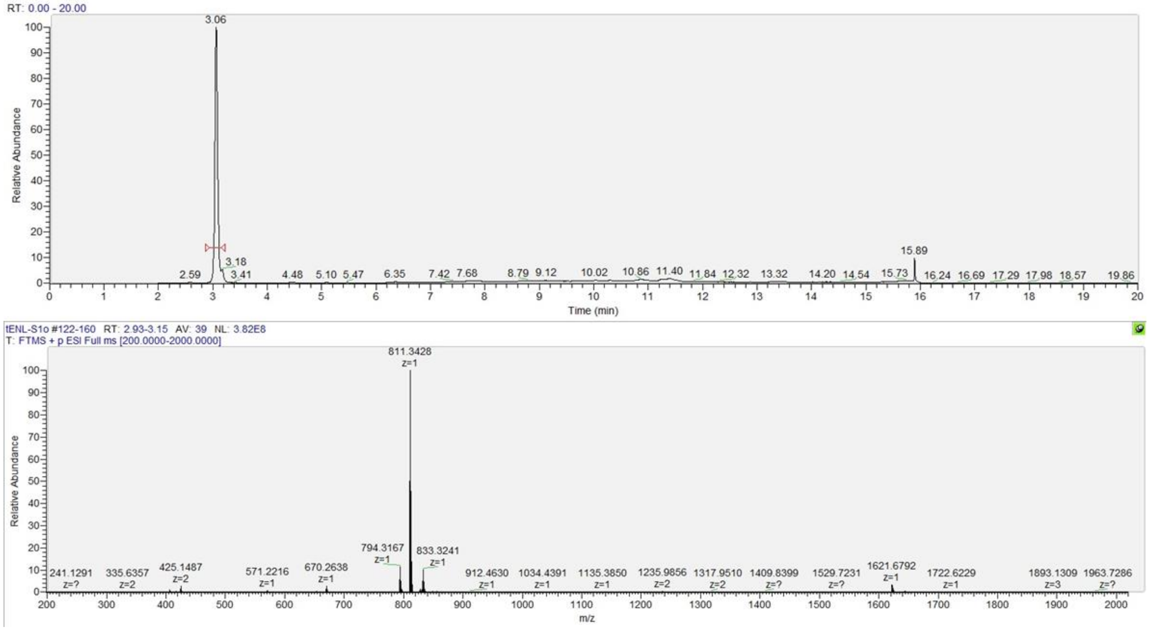


Figure S27. LC/MS of tENL-S1o. Calculated $[M+H]^+$: 811.3449 Da.
